# AutoWrinkleID: a machine learning pipeline for biofilm wrinkle identification and quantitative analysis

**DOI:** 10.64898/2026.09.16.751705

**Authors:** Mirko Murgia, Mo Vali, Abhirup Mookherjee, Nikhil Krishnan, Vincent Grumbacher, Jordan Romeyer-Dherbey, Diana Fusco

## Abstract

Biofilms are widely distributed in both natural and engineered systems and play a fundamental role in microbial ecology and biotechnological applications. When grown on agar substrates, biofilms often exhibit macroscopic structural features due to mechanical instabilities driven by matrix production. These wrinkles encode relevant information about biofilm growth and structural development and have even been shown to possess some functional roles. Despite their relevance, currently available approaches for wrinkle detection from images rely on manual annotation, which is time-consuming, not scalable, and strongly dependent on the annotator. In the present study, we developed an end-to-end machine learning pipeline, named AutoWrinkleID, that starts from bright field images of biofilms and performs wrinkle identification, characterisation, and quantification across multiple bacterial species and strains. The name reflects the automated identification and subsequent characterisation and quantification of biofilm wrinkles. The annotations used to generate the masks, namely images containing only the wrinkle structures, were obtained both through manual labelling and using constitutive fluorescence and motility fluorescence. Interestingly, we found that models trained using only manual annotations perform worse than those trained using motility fluorescence markers. A comparison between motility and constitutive fluorescence masks further indicates that motility based annotations consistently outperform constitutive fluorescence across all strains, strongly suggesting that motile phenotypes are tightly associated with wrinkle structures. Additional computational analysis is conducted on the same dataset to assess the robustness and limitations of the results as a function of training dataset.

We also observed that Sholl analysis, originally developed for the study of neuronal dendrites and later applied to wrinkle analysis, can effectively characterize wrinkle patterns, although its applicability is strongly limited by wrinkle shape and by the quality of the masks.

## 1 Introduction

Biofilms are ubiquitous in both natural and engineered environments and play a crucial role in microbial ecology and a wide range of biotechnological applications [1]. Among their most distinctive morphological features when grown on agar substrates is the formation of wrinkles, which can arise across different biofilm-forming species and exhibit a broad diversity of shapes and scales. In this study, the focus is placed on large-scale wrinkles that span significant portions of the biofilm structure [2]. The application of machine learning (ML) techniques in biofilm research is well established, both for integration with advanced imaging methods [1] and for the analysis of wrinkle patterns [3]. However, previous studies lack a robust and generalizable pipeline for the automated extraction of wrinkles directly from biofilm images, limiting their characterisation to time-consuming and bias-prone manual annotation.

In this work, we propose a complete machine learning pipeline, named AutoWrinkleID, in which a trained ML algorithm is capable of extracting wrinkle masks from biofilms formed by bacterial species different from those used during training, thus demonstrating a degree of flexibility and generalization in wrinkle identification. Furthermore, we show that the model can be effectively trained using masks that are not manually annotated, but instead derived directly from fluorescence signals of the biofilm formed by engineered strains, providing an alternative and scalable approach to ground truth generation. The use of two distinct fluorescence channels enabled the extraction of complementary information from the biofilm, specifically constitutive fluorescence and motility related fluorescence. These signals were exploited to identify wrinkle structures and to generate the corresponding masks, providing an alternative to manual annotation.

Multiple datasets were employed for both training and testing. For the training phase, strains of both *Bacillus subtilis* and *Escherichia coli* were used in order to enhance the flexibility of the identification algorithm. The testing phase also involved *B. subtilis* and *E. coli*, but with different strains and growth conditions to assess generalization capabilities. The datasets are summarized in Tab. 1.

For the task of identifying wrinkles in biofilm images, self supervised learning algorithms were adopted, following previous studies that demonstrated the suitability of this approach in scenarios where annotated data are scarce [1, 4]. In the context of this work, annotations correspond to the masks used to identify wrinkles in biofilm images. These masks may be generated either manually or through fluorescence based signals emitted by engineered bacterial strains. However, both strategies present limitations, since manual annotation is labour intensive, while fluorescence-based mask generation is not always easily transferable to all bacterial species.

Within this framework, the model learns directly from unlabelled data, which are subsequently incorporated into a fine-tuning stage that enables the extraction of the desired features. This approach has already been successfully applied to the analysis of cells and microbial by-products in biofilm images [4]. In line with previous studies that employed self-supervised learning for pixel-level classification in biofilm images [4], the behaviour of the model was further investigated by analysing its performance across different train/test splitting sizes in the same dataset.

Subsequently, the masks, and therefore the corresponding biofilms, were analysed using Sholl analysis. This method consists of drawing a series of concentric circles centred on the biofilm, allowing the study of wrinkle distribution within each annular region. This methodology has been employed for decades in the analysis of neuronal dendrites due to its ability to quantitatively describe their complex morphology [5]. Such structures exhibit similarities to biofilm wrinkles, as illustrated in Fig. 1.

**Fig. 1.**
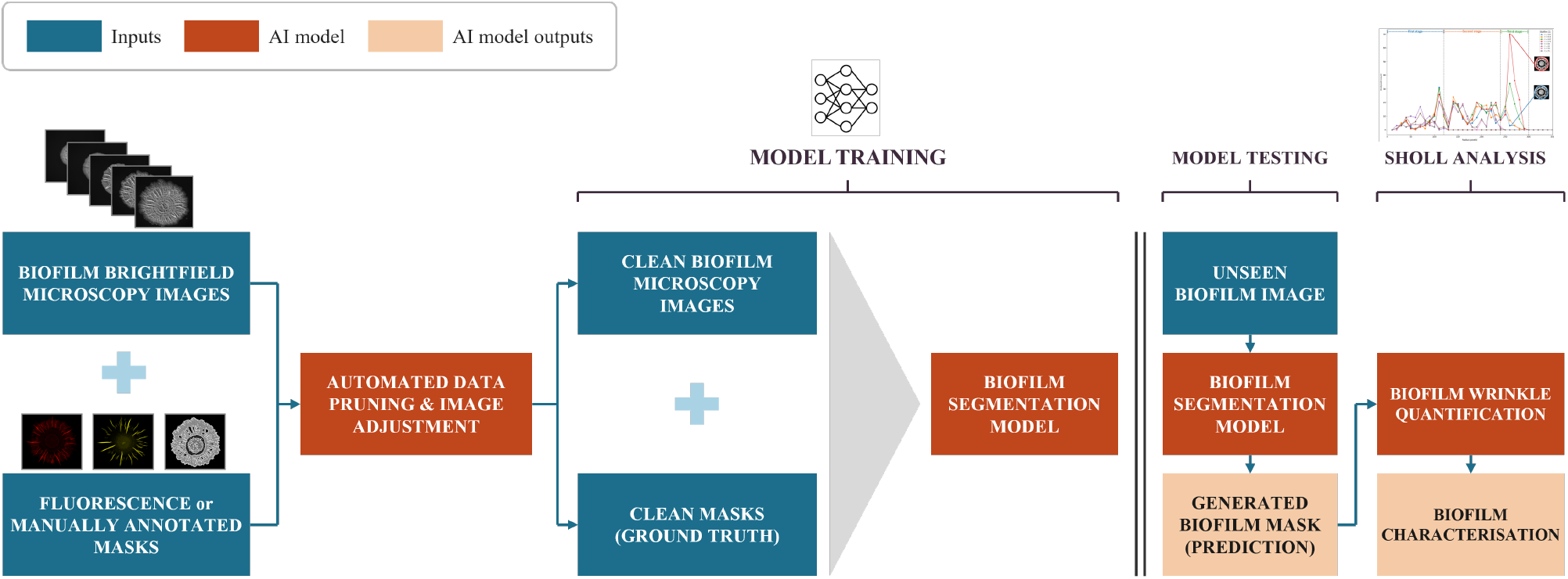
Overview of the biofilm wrinkle segmentation and Sholl analysis pipeline. The figure summarizes the entire pipeline: preprocessing, model training, model testing, and the final Sholl analysis. The aim is to develop a complete pipeline that, once trained on diverse biofilms and their corresponding masks, is able to extract and subsequently quantify wrinkle structures in previously unseen biofilms.

This methodology has already been successfully applied to the analysis of wrinkles in *B. subtilis* biofilms [3]; however, in the present study it is further extended and refined by introducing additional morphological descriptors, namely branching points and endpoints. These features improve the analysis of the well-established concept of biofilms in different developmental stages, by increasing the number of quantitative parameters available for biofilm characterisation. In this way, the proposed approach provides a more detailed description of biofilm morphology and development, going beyond the sole quantification of wrinkle spatial distribution. A schematic overview of the workflow is presented in Fig. 1.

### 1.1 Novelty of work

The present work introduces several methodological and experimental advancements over existing approaches through the proposed pipeline, AutoWrinkleID. A hybrid masking strategy is employed, combining manually annotated masks with fluorescence based masks derived from motility and constitutive fluorescence signals. This integration enhances segmentation robustness compared to previous studies that relied exclusively on manual annotation, and, more importantly, improves scalability. Indeed, this work demonstrate that reliable masks can be generated from fluorescence images and that, through the use of the proposed algorithm, valid wrinkle masks can be reconstructed even when only a limited number of annotated data are available. A rigorous data partitioning protocol is implemented, including clearly defined training, validation, and test sets. Unlike prior work, where validation and testing procedures are not explicitly described, this approach enables a reliable assessment of model generalizability beyond the training data. The dataset is expanded to include multiple bacterial strains and biofilm types during both training and testing phases. Earlier studies were restricted to a single biofilm type and growth condition, whereas the present work incorporates multiple bacterial species and strains grown in different conditions in both the training and testing phases, thereby enhancing the flexibility of the proposed algorithm and providing evidence of its scalability across species and culture conditions. In addition, a more detailed Sholl analysis is introduced, including additional descriptors such as branching points and endpoints. This enables a deeper characterisation of structural properties and improves the quantitative evaluation of biofilm morphology.

## 2 Materials and methods

### 2.1 Biofilm growth and image acquisition

Different strains of *B. subtilis* and *E. coli* were used for biofilm imaging (Table 1). *B. subtilis* strains were grown on MSgg agar [6], while most *E. coli* strains were grown on Salt-free LB (SFLB) agar [7]. MSgg plates were prepared one day prior to the experiment [6], left overnight at room temperature to solidify, and dried for an additional 1 h in a laminar flow hood immediately before inoculation. SFLB plates were prepared following standard procedure [7]. For inoculum preparation, all strains were grown overnight at 37 ^°^C in Luria-Bertani (LB, Invitrogen) medium with shaking at 250 rpm. For all biofilm experiments, 1 *µ*L of the respective inoculum was spotted onto the surface of the agar plates. After inoculation, MSgg plates were sealed with masking tape, and SFLB plates were sealed with double-layer of parafilm. Plates were incubated at their respective optimal temperatures (Table 1). Specifically, *E. coli* strains ATCC 25922 [8] and CFT 073 [9] were grown on synthetic urine medium supplemented with 1.5% agar and incubated at 37 ^°^C for six days prior to snapshot imaging. Imaging was carried out using a Carl Zeiss Axio Zoom.V16 stereo microscope equipped with a Zeiss PlanApo Z 0.5×/0.125 FWD 114 mm objective lens. Images were captured either as single time-point snapshots or via time-lapse imaging (40 min intervals). An integrated incubation chamber maintained the required growth temperatures during time-lapse imaging acquisition (Table 1). For each experiment, bright-field images were acquired to document overall colony morphology and wrinkle architecture. Fluorescence images were acquired using corresponding filter channels (mCherry and mScarlet: 587/610 nm; mTurquoise: 433/475 nm; YFP: 508/524 nm) to generate fluorescence-derived wrinkle masks, which were subsequently compared with manually annotated reference masks. In *B. subtilis*, the P*hag*, P*tasA*, and P*veg* promoters were used as reporters for motility, matrix production, and constitutive gene expression, respectively. In *E. coli*, the P*fliC*, P*csgD*, and P*gapA1* promoters served equivalent respective functions. The Manual dataset consists of a *B. subtilis* NCIB 3610 biofilm expressing both flagellar and matrix reporters, grown on MSgg agar (1.5%) at 30 ^°^C. This dataset includes biofilms ranging from 3 to 5 days old. The Constitutive I and Motility I datasets originate from *E. coli* strain AR 3110 expressing constitutive, flagellar, and matrix reporters. These were incubated at 28 ^°^C on SFLB medium supplemented with varying agar concentrations (1%, 1.5%, and 2%). The Constitutive II and Motility II datasets utilize two different *B. subtilis* strains (either expressing only constitutively or constitutive plus-flagellar motility) grown at 30 ^°^C. Nutrient conditions consisted of MSgg medium containing 1× glycerol and varying concentrations of glutamate (0.5×, 1×, and 1.5×). The Test 1 dataset was generated from a competition experiment. Overnight cultures of non-flagellated (Δ*hag* -KmR; amyE::P*vegS* -mScarlet-specR) and non-matrix (Δ*epsA-O* -TetR; amyE::P*vegS* -mTurquoise-specR) *B. subtilis* strains were washed three times with phosphate-buffered saline (PBS). Cell suspensions were adjusted to an OD_600_ of 1.0 and mixed at a 1:1 ratio to generate the competitive inoculum [6]. The Test 2 dataset was derived from *B. subtilis* strain 3a38 (expressing constitutive and flagellar genes) grown at 30 ^°^C on MSgg containing 0.3–1× glycerol and 0.5–1× glutamate. The Test 3 dataset comprises *E. coli* snapshot images acquired under distinct growth conditions. Strain 536 [10] was cultured on SFLB agar, while strains ATCC 25922 [8] and CFT 073 [9] were cultured on synthetic urine agar and imaged after six days of incubation at 37 ^°^C. Crucially, these variations in bacterial strains, growth media, and nutrient concentrations were intentionally selected to induce a wide spectrum of structural complexity, generating distinct biofilm morphologies ranging from robust, well-defined wrinkles to poorly developed or atypical architectures. Collectively, this imaging workflow produced spatially matched bright-field and fluorescence datasets across varied bacterial strains, nutrient conditions, and developmental stages (Table 1), providing the input images and ground-truth masks necessary for subsequent machine learning-based wrinkle identification.

**Table 1.**
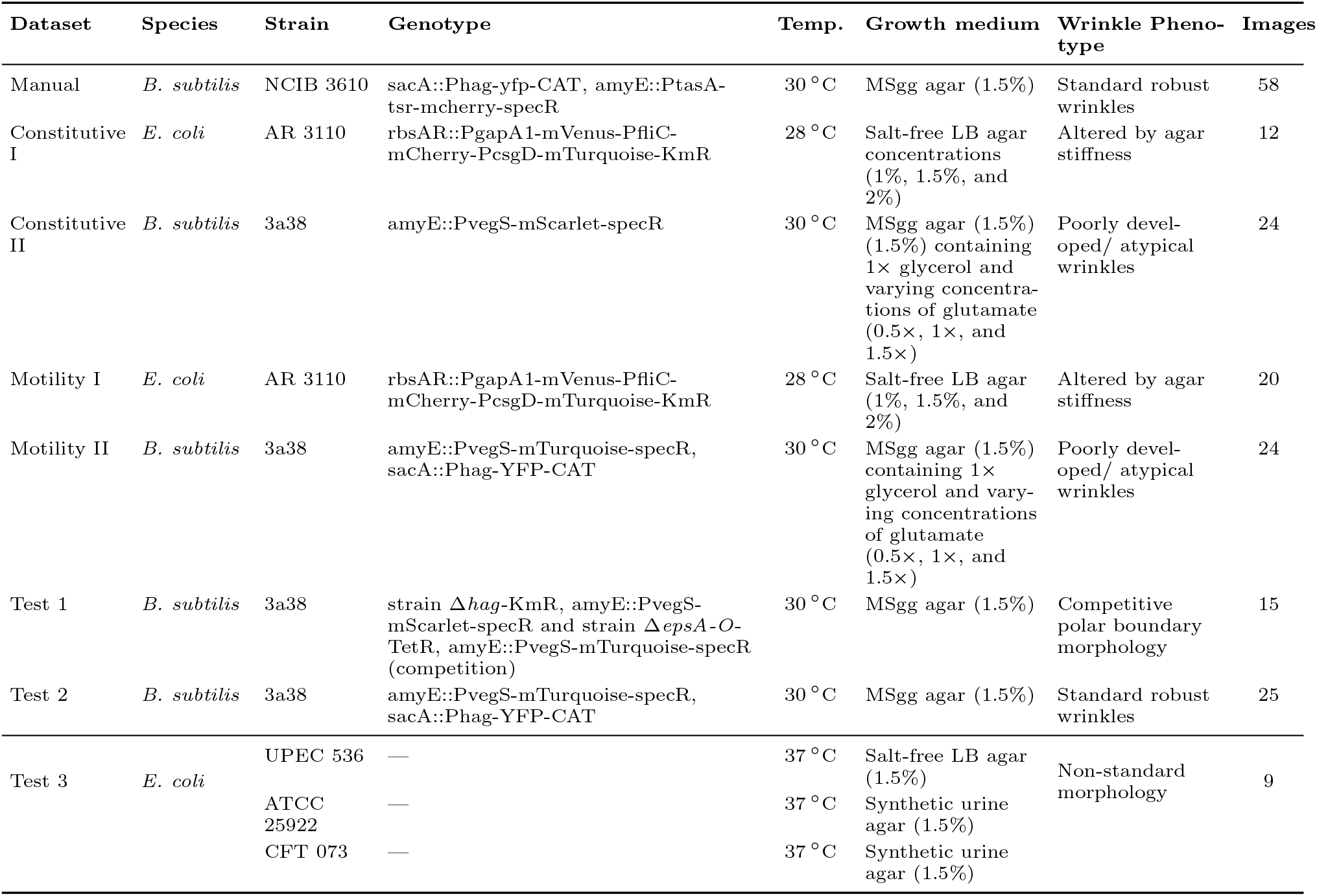
Summary of bacterial datasets. The table reports the bacterial species, strains, genotypes, incubation temperatures, growth media, and number of images employed for training and testing. Additional details on growth conditions and image acquisition are provided in the subsection entitled “Biofilm Growth and Image Acquisition”.

### 2.2 Image preprocessing description

Because one biofilm typically exhibits multiple independent wrinkles, to address the limitation of acquiring a large biofilm dataset, without relying on data augmentation techniques, a patching strategy was adopted. Both images and their corresponding masks were subdivided into smaller regions. This approach provides a twofold advantage over data augmentation. First, it increases the number of available training samples, allowing the model to be trained on a larger dataset. Second, each patch represents a localized portion of the biofilm, thereby reducing the complexity of individual inputs compared to full biofilm images. The dataset consists of images extracted from multiple time-lapse sequences. A careful selection of images was performed prior to training, excluding biofilms that did not exhibit wrinkle formation, the initial frames of each time-lapse sequence in which wrinkles had not yet developed, and images captured within short temporal intervals (approximately 20/30 frames), as these do not present significant structural differences compared to preceding frames. Following this selection process, both images and corresponding masks were preprocessed using the FIJI distribution (Version 1.54m) of *ImageJ* [11] in order to adjust brightness and contrast, thereby providing the model with the best possible input quality. Subsequently, the masks were binarized, as the model operates in a binary classification framework: each pixel is classified either as belonging to a wrinkle (white) or as background (black). This process is schematically illustrated in Fig. 2.

**Fig. 2.**
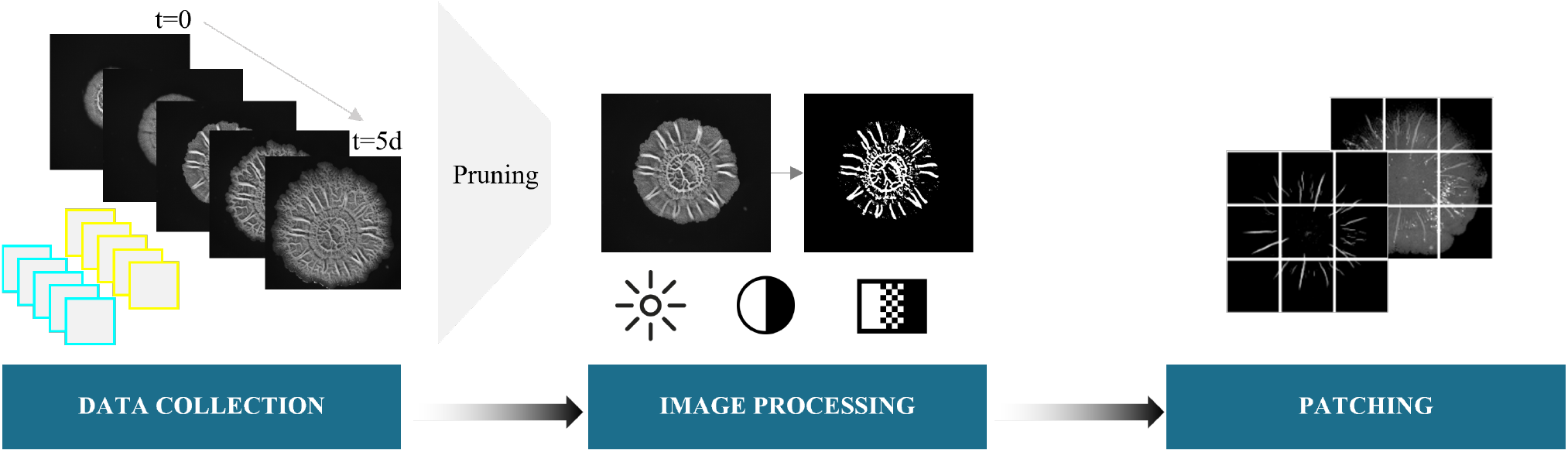
Overview of the image processing pipeline. Data are collected from time-lapse series of microscope images of biofilms, then adjusted for brightness, contrast and greyscale, before being patched into small localised portions of the biofilm image.

### 2.3 Creation of ground truth masks

The ground truth fluorescence masks were generated using genetically modified bacterial strains carrying fluorescent reporters, as described above. Images were acquired through different fluorescence channels of the same microscope. The constitutive reporter was used to label the overall bacterial population and therefore to delineate the structure of the biofilm, whereas the motility reporter produced a fluorescence signal associated with the expression of motility related genes, allowing the spatial distribution of motile cells within the biofilm to be visualized. As described in the previous section, mask generation included a preprocessing step aimed at enhancing the contrast between the fluorescence signal associated with the wrinkles and the surrounding biofilm. Brightness, contrast, and greyscale levels were adjusted for each image individually, using different threshold values selected to make the wrinkles as clearly distinguishable as possible from the rest of the biofilm.

The manually annotated masks, corresponding to the Manual dataset, were created by members of the research group with previous experience in biofilm analysis. A standard image editing program was used to place an empty annotation layer over the corresponding brightfield image. The visible wrinkles were then manually traced, taking into account both their morphology and thickness. The resulting annotation mask was superimposed on the original brightfield image to verify that the annotated wrinkles accurately overlapped the structures visible in the image, including their spatial extent and thickness.

### 2.4 Model architecture

The Barlow Twins (BT) method, inspired by the redundancy reduction principle proposed by neuroscientist H. Barlow, is a self-supervised learning algorithm [12]. The method measures the cross-correlation matrix between the outputs of two identical neural networks, the twins, fed with distorted versions of the same sample, with the objective of approximating the identity matrix. This minimises redundancy and improves robustness of the learned representations [12].

Our implementation of the algorithm was inspired by a publicly available code repository [13]. The BT method is fed with two distorted version of the same image and the output is passed to a segmentation head, which generates the predicted mask. The predicted mask is compared with the ground truth mask, allowing the segmentation module to learn the desired wrinkle extraction task. This algorithm, typically employed for the classification of regions or entire images, is here adapted to perform pixel-wise classification, producing a generated mask. The availability of a ground truth mask enables the evaluation of the similarity between the predicted and the reference mask at each stage of the algorithm. The training process is therefore driven by the minimization of two loss functions: the first associated with the cross-correlation matrix, and the second related to the discrepancy between the generated mask and the ground truth mask.

By reducing redundancy, the BT approach aims to generate embeddings that are invariant to distortions and statistically independent across different parts of an image. This ensures that neural networks trained with this method produce representations capable of capturing the essential features of the data, while discarding redundant information [12]. This property makes the Barlow Twins algorithm particularly robust and well suited to the identification of wrinkles in biofilm images. Moreover, this algorithm has already been used in the literature for the identification of bacterial species in microscopic images [4]. The same study also reported the use of another self supervised learning algorithm, MoCoV2, which was also tested during the initial stages of the present work. However, MoCoV2 was subsequently discarded because it showed lower performance in terms of the selected evaluation metrics, namely accuracy and Dice score.

### 2.5 Sholl analysis

Sholl analysis is performed on the binarized masks containing only the wrinkle structures of the biofilm [5]. The method further relies on a ”skeletonisation” step, which is essential for quantifying the spatial distribution of the structure with respect to the concentric circles. The skeleton was generated using the *skeletonize* function from the *skimage*.*morphology* module of the Python *scikit image* library and consists of a one pixel wide representation of the wrinkles derived from the binary mask [14]. The centre of the biofilm is subsequently identified through an automatic procedure based on image intensity distribution. A series of concentric circles is then constructed, with a fixed radial increment of 10 pixels, a value selected as appropriate for capturing the structural complexity of the analysed dataset. All parameters are evaluated as a function of the radial distance from the centre, expressed in pixels. The investigated metrics include the number of skeleton pixels within each annular region, providing a measure of wrinkle distribution, the corresponding pixel density per annulus, and topological descriptors such as the number of branching points, endpoints, and internal points. A branching point is defined as a location where the skeleton splits into two or more segments, an endpoint as a terminal point of the skeleton, and an internal point as any point that is neither a branching point nor an endpoint. The disk count for annulus represents the number of skeleton pixels within successive radial regions, obtained by considering the incremental difference between concentric disks. This parameter provides a radial distribution profile of the biofilm structure, rather than a direct count of intersections as defined in classical Sholl analysis.

## 3 Results

Several datasets composed of images and corresponding masks were used for training after preprocessing. In addition to individual datasets, different combinations were considered to improve model robustness and generalisability. Training was performed separately using masks derived from constitutive fluorescence and masks derived from motility fluorescence. Each fluorescence based dataset was evaluated both independently and in combination with the manually annotated mask dataset. The performance of each configuration was evaluated using both quantitative metrics and qualitative inspection of the validation outputs. The weights obtained from the best performing configurations were then selected and used for testing on datasets without annotated masks. Finally, Sholl analysis was performed on the Manual dataset. This choice was motivated by the fact that this dataset provides the most reliable reference masks for the quantitative analysis. In these images, wrinkles are more clearly distinguishable from the biofilm background than in the fluorescence derived datasets, and the masks do not contain the small segmentation anomalies that may appear in the generated testing masks. Therefore, the Manual dataset was selected to establish and validate the Sholl analysis procedure under the most controlled conditions, using the highest quality masks available. This avoids propagating possible segmentation artefacts into the downstream morphological characterisation.

### 3.1 Segmentation

Each dataset was randomly split into a training set and a validation set, using an 85/15 ratio. The validation dataset, unlike the training dataset used to fit the model, provides an unbiased estimate of the model’s ability to generalize to new, unseen data. This subset plays a crucial role both in the optimization of the model parameters, such as the network weights, and in the tuning of hyperparameters. For this reason, the results presented, in terms of both evaluation metrics and generated outputs, refer to the validation subset of each training dataset, as they provide a more representative and informative assessment of the model’s learning process compared to the training data.

The hyperparameters evaluated in this study include the learning rate and the batch size. Model performance was assessed using the loss function, accuracy, and the Dice score [15, 16]. The loss function reported refers to the segmentation component, namely the comparison between the generated mask and the ground truth mask, as it is more directly related to the final output of the model and therefore more representative of its performance. In contrast, the loss function associated with the Barlow Twins framework, based on the cross-correlation matrix, is primarily related to the representation learning process and the way the Barlow Twins extracts features from the input images. As such, it is not directly linked to the quality of the final segmentation output. All models were trained for 50 epochs, as convergence of the loss function was consistently observed at this stage. For hyperparameters tuning, the *Weights and Biases* online tool [17] was employed, as it enables real-time monitoring of metrics across different runs of the same training process. The analysis conducted on the various datasets yielded consistent results in the selection of the learning rate and batch size, as the datasets are comparable in size and exhibit relatively homogeneous characteristics. Of particular relevance, the optimal hyperparameters among those tested were a batch size of 32 and a learning rate of 10^*−*4^ across all datasets and their combinations. Although some differences are observed, the overall trends of the evaluation metrics remain relatively consistent. The datasets, their combinations, and the corresponding evaluation metrics are summarized in Tab. 2, while the evaluation curves are presented in the SI.

The evaluation curves, including loss, accuracy, and Dice score, exhibit a generally homogeneous behaviour across the different training configurations. In most cases, a consistent trend is observed between models trained on masks generated from motility fluorescence and those derived from constitutive fluorescence. The main differences between these approaches, namely whether the images used to generate the masks were acquired through motility fluorescence or constitutive fluorescence, do not emerge at this stage, but rather during the testing phase, where models trained on motility based masks demonstrate greater flexibility and improved accuracy. Overall, all curves show satisfactory performance, supporting the selection and saving of model weights for subsequent testing. However, only the most relevant models were retained, based on both validation curves and qualitative assessment of the generated outputs. In total, the weights from four datasets were selected.

Two of the selected models correspond to those achieving the highest Dice scores, namely Manual + Constitutive I and Manual + Motility I. These configurations present intermediate values in terms accuracy. The remaining two selected models correspond to Manual + Constitutive I + Constitutive II and Manual + Motility I + Mo1 combinations including all datasets within each category. They show balanced performance in terms of accuracy and relatively strong Dice scores. The weights obtained from these training configurations were selected due to their superior flexibility in generating accurate and generalizable segmentation masks, as shown in the next subsection.

While the analysis of evaluation metrics is essential, the qualitative assessment of the model outputs is equally important. It is therefore crucial to examine not only the differences between the generated masks and the ground truth, but also their consistency with the original images from which the wrinkle structures are extracted. In several cases, the algorithm demonstrates a superior ability to identify wrinkle structures compared to manual annotations, an example in Fig. 3, likely because it learns directly from the original images in addition to the provided masks. However, limitations in wrinkle identification are more evident in specific conditions, particularly in datasets fluorescence-based datasets of *B. subtilis* (C2 and Mo2), where variability in the signal and dataset boundaries may affect the accuracy of the segmentation, as shown in SI.

**Fig. 3.**
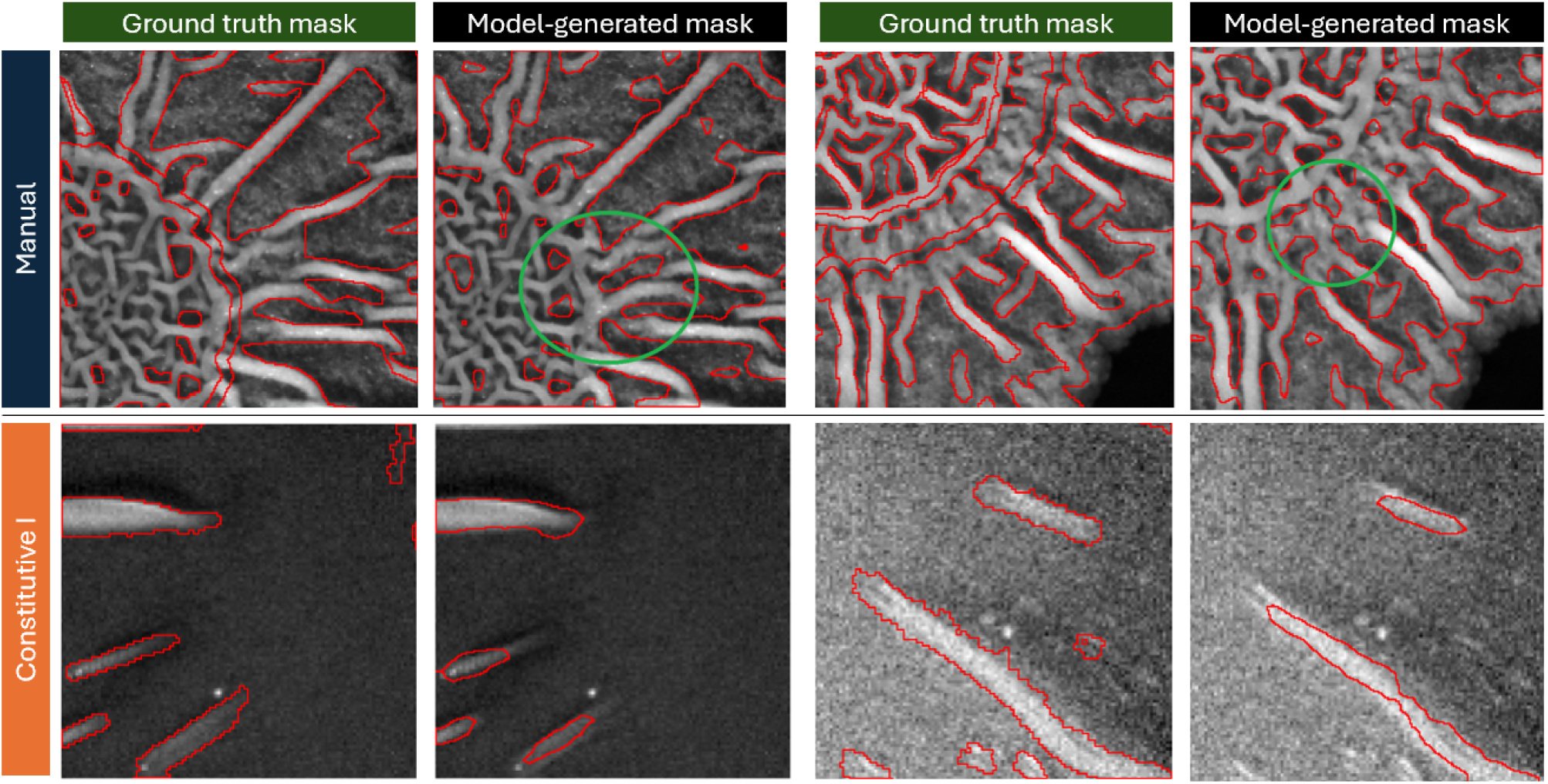
Outputs from model trained on M+C1 datasets. These examples are considered representative of the training behaviour for this dataset, the images come from validation dataset, whose weights were selected for the testing phase. In the first row, the green circle highlights how the model captures the wrinkle pattern more accurately than the manual annotation shown, particularly in the outer regions. The model also performs well on the second row, despite the wrinkle pattern being simpler, because the contrast between wrinkles and the surrounding biofilm is lower, making segmentation more challenging.

The results of the training phase are presented below for the datasets considered most significant in terms of both evaluation metrics and qualitative outputs. As for the evaluation metrics, the reported results are derived from the validation subsets, as they provide a more reliable assessment of model performance. The datasets included in this analysis are M+C1 and its motility-based counterpart, M+Mo1, as well as TotC and TotM. For convenience, these latter combinations are referred to as TotC and TotM, respectively.

#### 3.1.1 M+C1 and M+Mo1 datasets

After the model is trained, we validate the model using unseen images from the held-out datasets of M+C1 and M+Mo1, the analysis of which is shown in Fig. 3 and Fig. 4, respectively. The original patched image is displayed with the ground truth mask overlaid on the left and the mask generated by the model on the right. In both figures, the results show that for patches originating from the M dataset, corresponding to the first row of each figure, the model performs better than the manual annotation by capturing additional details, particularly in the outer regions. This behaviour is related to the use of Barlow Twins, a self supervised learning approach, which learns directly from the original images, while the comparison with the annotation masks is introduced only in a subsequent stage.

**Fig. 4.**
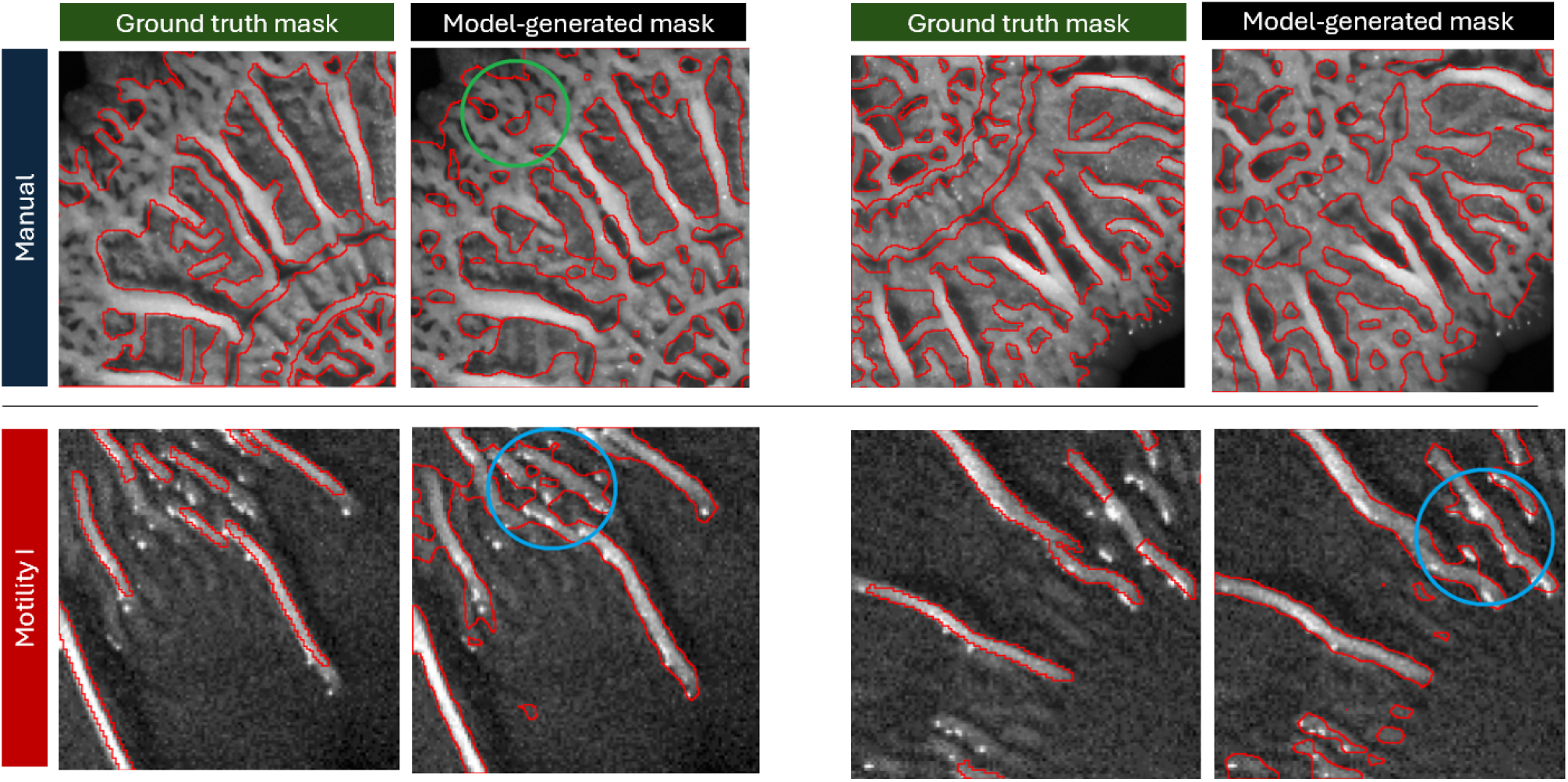
Outputs from model trained on M and Mo1 datasets. These examples illustrate the effective performance of the model when trained on this dataset. The green circle in the second panel in the first row highlights how the model detect the complexity of the wrinkle pattern more accurately than the manual annotation. The blue circles in the panels in the second row indicate two cases in which the model delineates wrinkle structures more precisely than the motility fluorescence based masks.

The model shows slightly greater difficulty, while still providing correct identification, for data originating from C1 and Mo1, corresponding to the second row of the figures. Although the wrinkles in the M dataset of *B. subtilis* exhibit greater structural complexity than those observed in the *E. coli* datasets C1 and Mo1, the model achieves better segmentation performance on the M dataset. A possible factor contributing to this result is the clearer distinction between the wrinkles and the surrounding biofilm in the M dataset compared with C1 and Mo1. Therefore, model learning appears to be influenced not only by wrinkle morphology, but also by how distinctly the wrinkles stand out from the surrounding biofilm.

#### 3.1.2 TotC and TotM datasets

In the following section, outputs from the training phase of the model on the TotC and TotM datasets are presented. Once again, the displayed results correspond to the validation subsets, as they are considered the most representative for assessing the effectiveness of the training process. Fig. 5 presents representative patches from the TotC training, whereas Fig. 6 shows representative examples from TotM. In both figures, the first row includes images originating from M. Also in this case, the model produces masks that delineate wrinkles more precisely than the manual annotations. The second row of both figures shows results from the C1 and Mo1 datasets, respectively. In both cases, the model appears more transferable in situations where wrinkles are more difficult to distinguish from the biofilm background. The bottom row presents representative patches illustrating how the Barlow Twins learns from data originating from C2 and Mo2. For the C2 samples, shown in Fig. 5, the model exhibits behaviour similar to that observed for C1, correctly identifying the wrinkle structures.

**Fig. 5.**
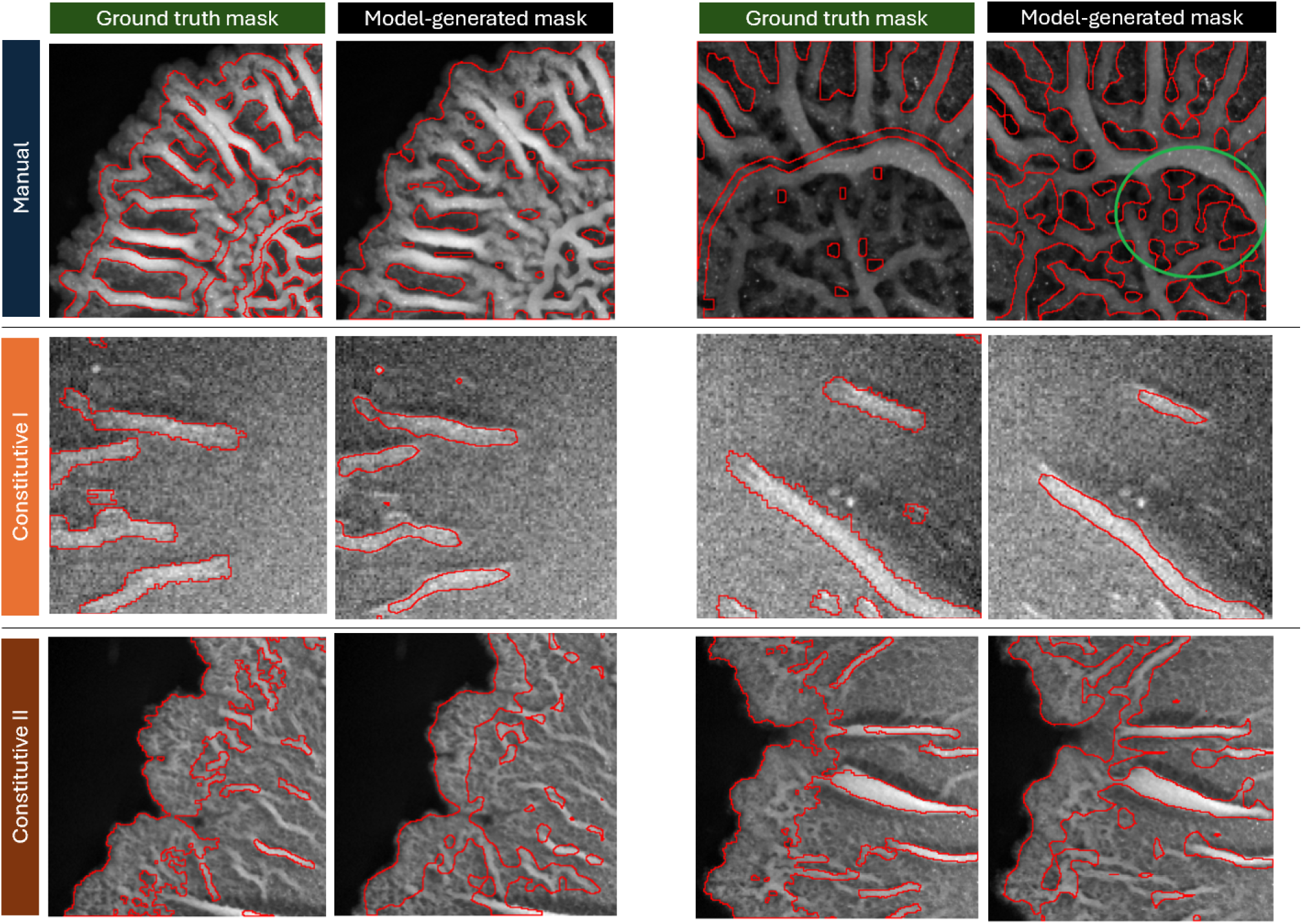
Outputs from model trained on TotC dataset. As in the previous cases, the model learns the wrinkle structures more accurately than the manual annotations. The green circle in the second panel in the first row highlights the complex central wrinkle region of the biofilm, which is effectively captured by the model. In the second row, corresponding to C1, the typical difficulty due to the low contrast between wrinkles and the surrounding biofilm is observed. In the third row, corresponding to C2, the model correctly captures the structure of larger wrinkles, while showing greater difficulty in identifying smaller wrinkles located in the outer regions of the biofilm.

**Fig. 6.**
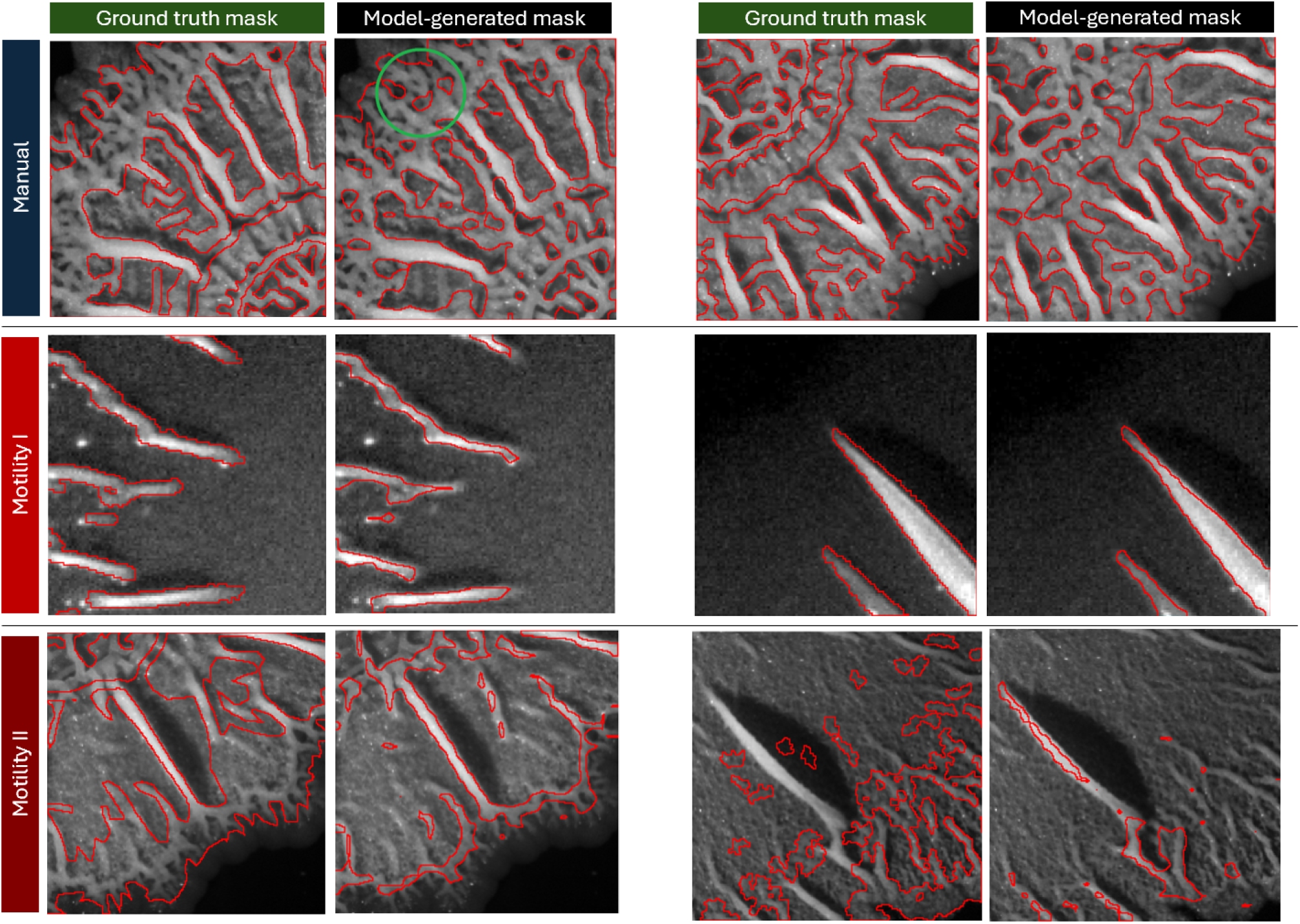
Outputs from model trained on TotM dataset. As in the previous cases, the first and second rows, corresponding to M and Mo1, respectively, show that the model learns effectively in these configurations. Greater difficulty is observed in the more complex scenario represented by the third row, corresponding to Mo2, where wrinkles are more numerous and intricate, and the contrast between wrinkles and the biofilm background is reduced.

As shown in Fig. 6, the TotM model is more reflective of the ground truth for the validation set containing the M dataset, than it is for the Mo1 and, particularly, Mo2 datasets. We posit this may be due to the well-established nature of wrinkles in the M dataset, comprising *B. subtilis* NCIB 3610 vs. the Mo1 (*E. coli* AR3110) and Mo2 (*B*.*subtilis* YFP CFP) datasets.

### 3.2 Performance on different biofilm morphologies

The objective of this section is to evaluate the performance of the model using different sets of weights for identifying wrinkles across different biofilm morphologies. Importantly, we want to assess the algorithm’s performance when trained on constitutive datasets versus its counterpart trained on motility datasets.

Looking purely at the model-generated outputs first, we find that the TotC model is able to generate the most visually correct masks overall (fourth column in Fig. 7) . However, the best performance in terms of validation accuracy is obtained with the TotM model (last column in Fig. 7). TotM gave 0.85 validation accuracy and 0.73 dice, whilst TotC gave 0.81 and 0.76, respectively. The higher accuracy of the TotM model can be illustrated in more difficult, low contrast scenarios where the biofilm is not clearly distinguishable from the background, such as the biofilm shown in the third row of Fig. 7. Although the TotC model generates a reasonably good quality mask for this biofilm, it misses a significant portion of wrinkles, in contrast to the TotM model which seems to generate more visually correct images. In these low contrast conditions, other models such as the M+C1 model, fail to capture the wrinkle structure entirely.

**Fig. 7.**
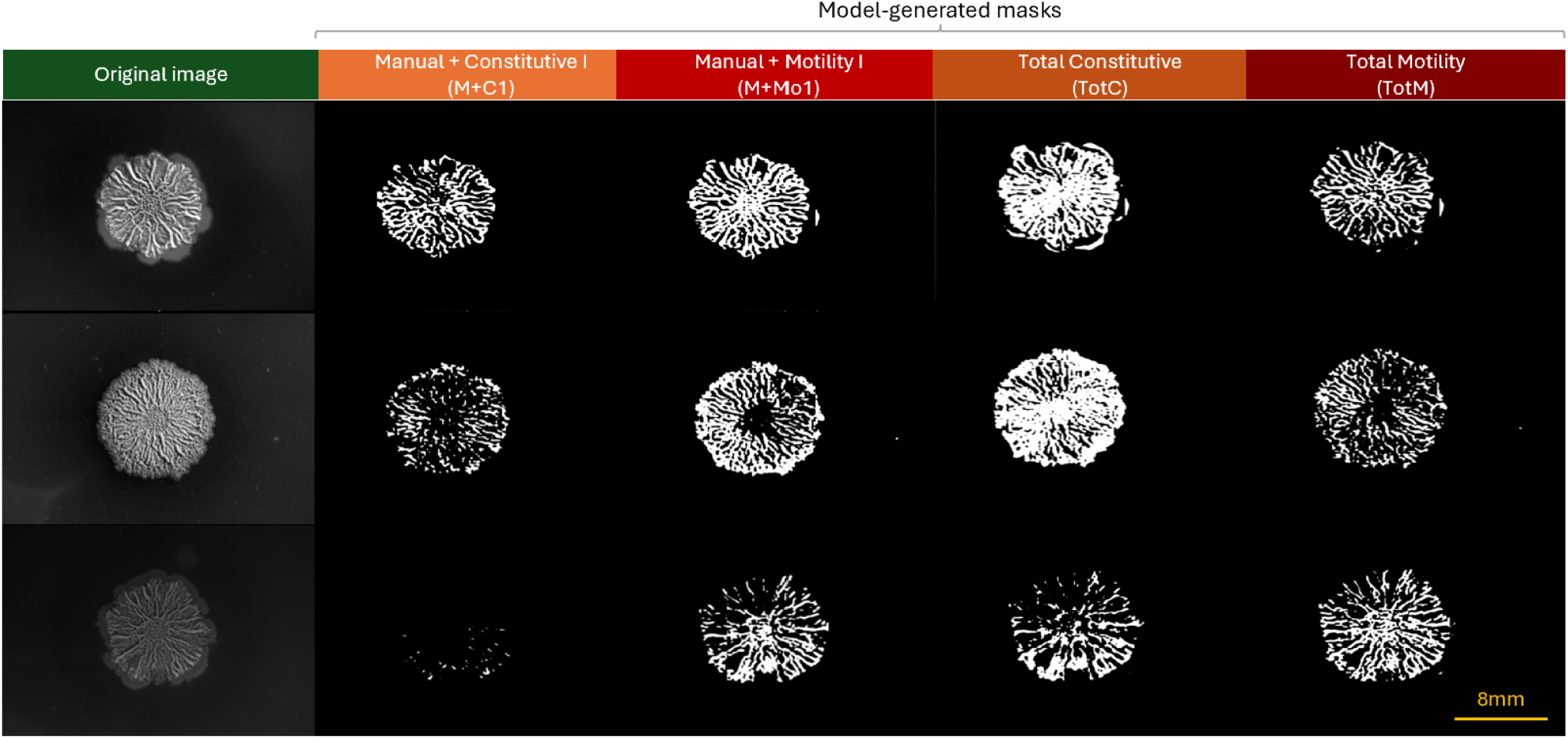
Results on Test 1. In all three examples, the masks generated using motility based weights, namely the last panel of each subgroup, represent the wrinkle patterns more accurately than the corresponding constitutive fluorescence predictions, shown in the second panel of each subgroup. The biofilms in this dataset are morphologically similar to those included in the M dataset. However, the improved performance obtained with the Total configurations, for both motility and constitutive fluorescence, indicates that training based solely on manual annotations is not sufficient to achieve consistent and robust wrinkle identification. These biofilms have diameters ranging from approximately 6 to 10 mm.

The algorithm was then tested on a *B. subtilis* dataset, referred to as Test 2, containing biofilms similar to those included in C2 and Mo2. This dataset, shown in Fig. 8, includes biofilms at different developmental stages. The configurations trained with TotM show greater adaptability in more complex scenarios, particularly in the last two images, and improved stability in simpler cases, corresponding to the first two images. The difficulty in the first two images arises from the presence of very small wrinkles with intensity values comparable to those of the more prominent wrinkles, namely those correctly identified by the algorithm. Despite the TotM model performing well in this scenario, small structures are still occasionally detected as wrinkles.

**Fig. 8.**
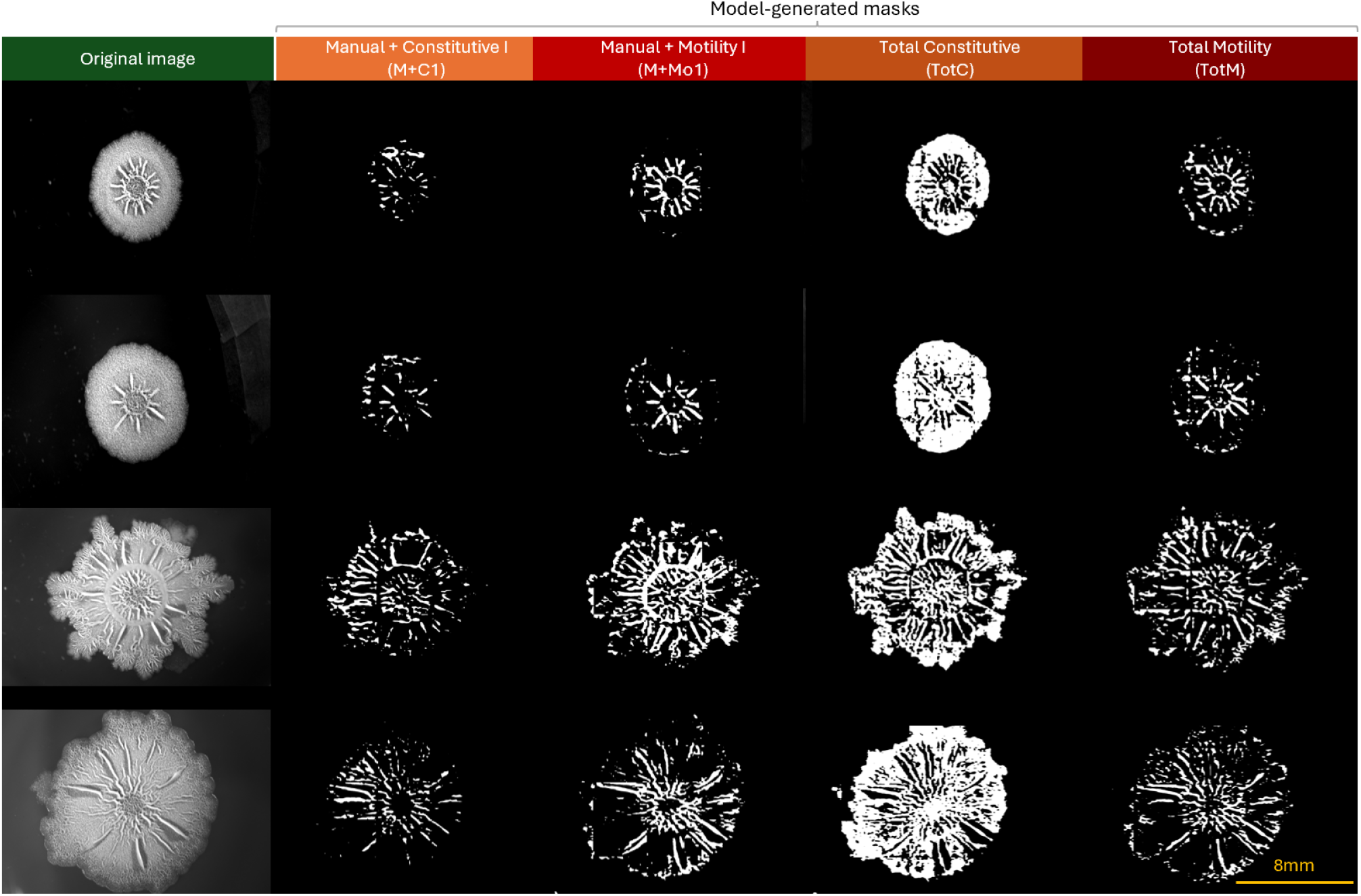
Results on Test 2. This test dataset, referred to as Test 2, includes two different types of biofilms: younger biofilms, aged one to two days, shown in the first and second rows, and older biofilms, aged ten days, shown in the third and fourth rows. The data in this dataset are similar to those in the C2 and Mo2 datasets. The segmentations obtained using the model trained on M+Mo1 panels, and those obtained using the model trained on TotM, panelS show accurate identification of wrinkle structures.In contrast, the corresponding models trained using constitutive fluorescence fail to capture the complexity of the older biofilms and also struggle to identify wrinkles in the younger ones, where the wrinkle luminance is very similar to that of the surrounding biofilm. These biofilms have diameters ranging from approximately 6 to 10 mm.

The model was further tested under a particularly challenging scenario, referred to as Test 3 in Fig. 9, which consists of a dataset of biofilm images exhibiting wrinkle patterns different from those observed during training, as they arise from uro-pathogenic *E. coli* strains. Specifically, these biofilms display thinner and denser wrinkles compared to the training data, and some samples do not present well defined wrinkles but rather blister-like structures. The configurations trained with TotC and TotM are still able to recognize and identify the general wrinkle pattern, although with lower accuracy than in Test 1 and Test 2. Remarkably, all configurations are able to identify the presence of blisters in bottom row, even though no similar biofilm patterns were included during the training phase.

**Fig. 9.**
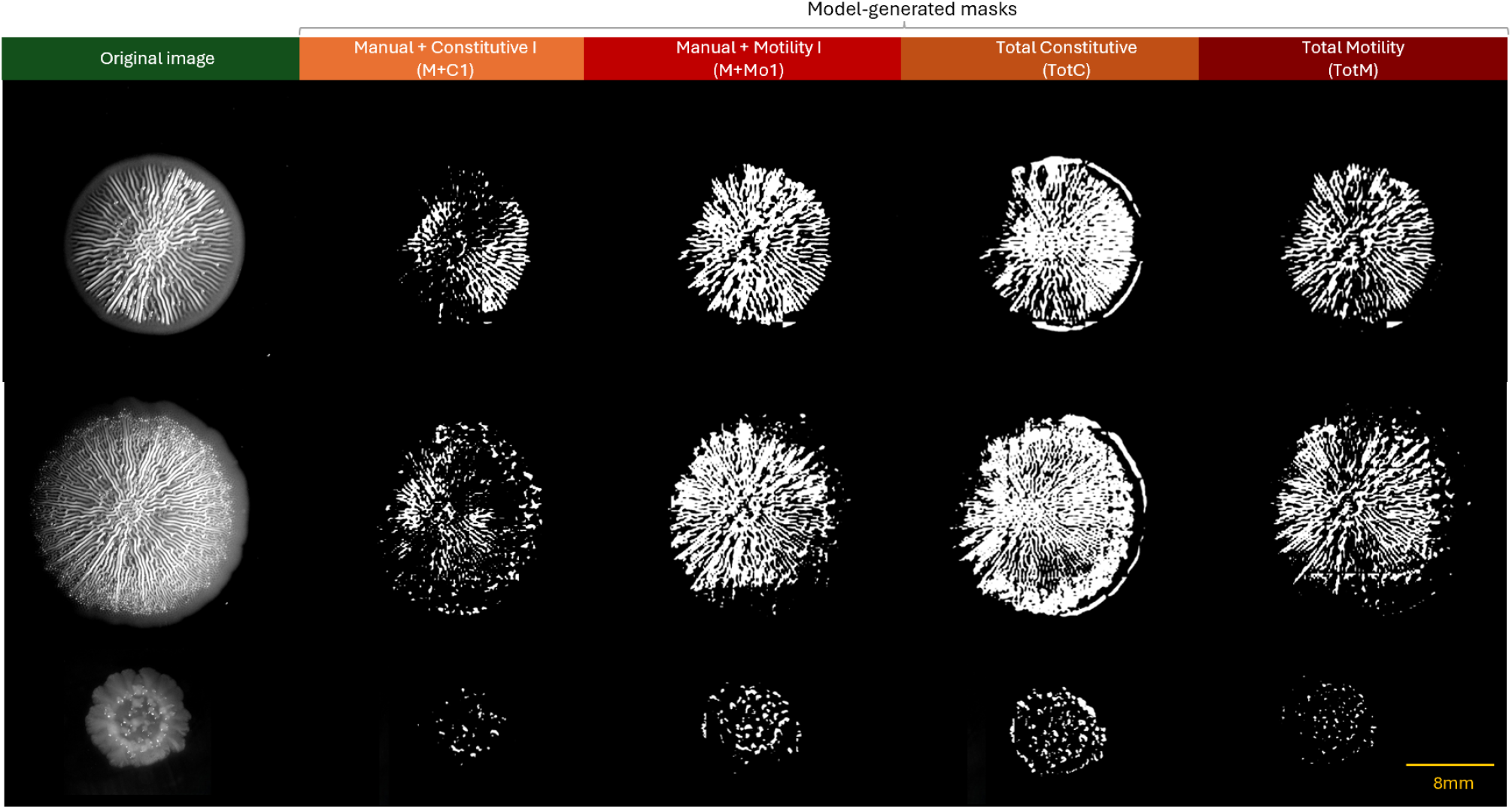
Results on Test 3. This test dataset, referred to as Test 3, represents a challenging scenario for the model, as the data differ significantly from those used during training. The dataset includes biofilms exhibiting fingerprint like wrinkle patterns, shown in the first and second rows, as well as biofilms that do not display classical wrinkles but rather bubble like structures. All configurations are able to identify both the fingerprint like wrinkles and the bubble structures. As observed in the previous examples, the models trained using motility fluorescence data show better performance than their counterparts trained using constitutive fluorescence.

The testing phase showed that the model is not only able to identify wrinkles under conditions similar to those used during training, but is also sufficiently flexible to adapt to more challenging scenarios. The weights obtained from training on the TotM dataset provided the best overall performance on the test images. Although this model does not achieve the highest quantitative metrics, as shown in Table 2, it performs best in terms of wrinkle identification on unseen test images that do not belong to the training dataset.

**Table 2.**
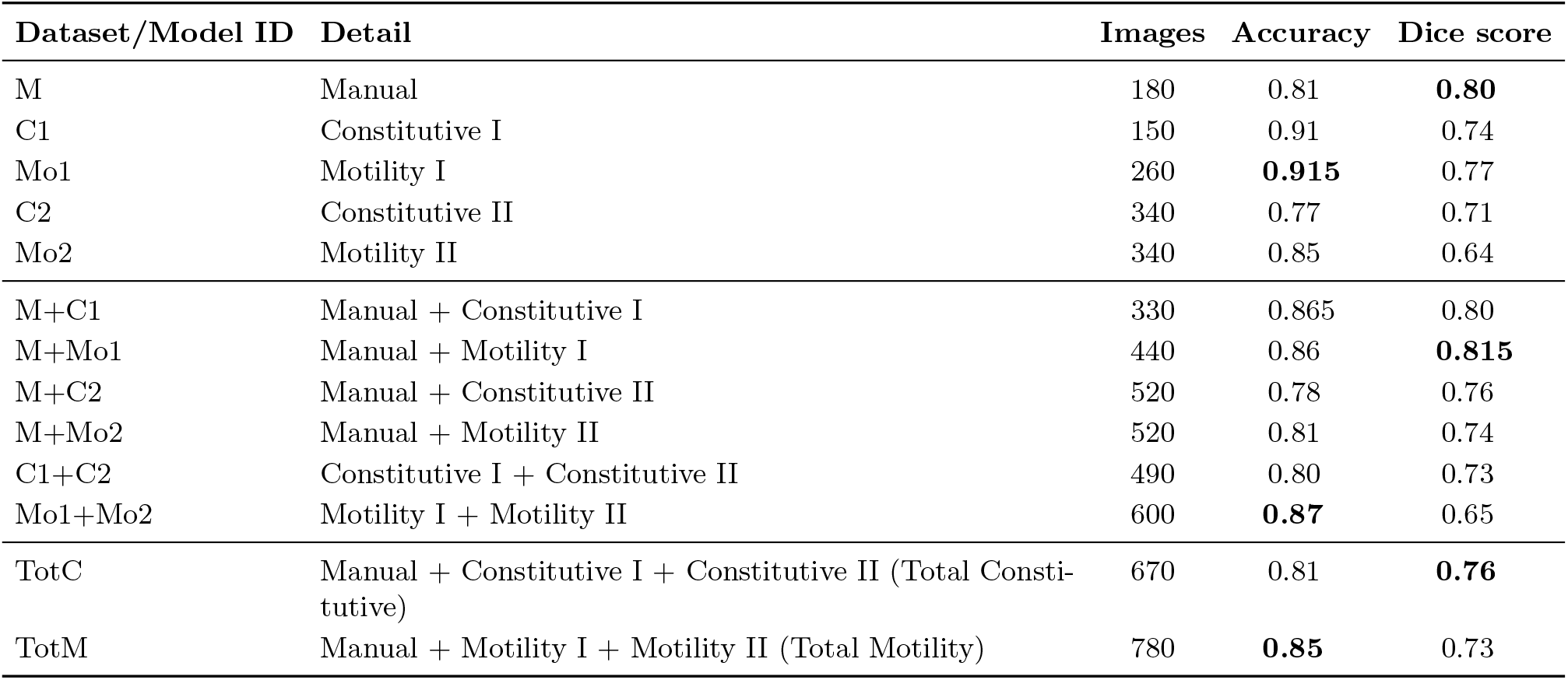
Summary of the datasets. Overview of the datasets used for training, including the number of patches and evaluation metrics. The learning rate was set to 10^*−*4^ and the batch size to 32. Results are reported for the validation subsets of each dataset and their combinations.

**Table 3.**
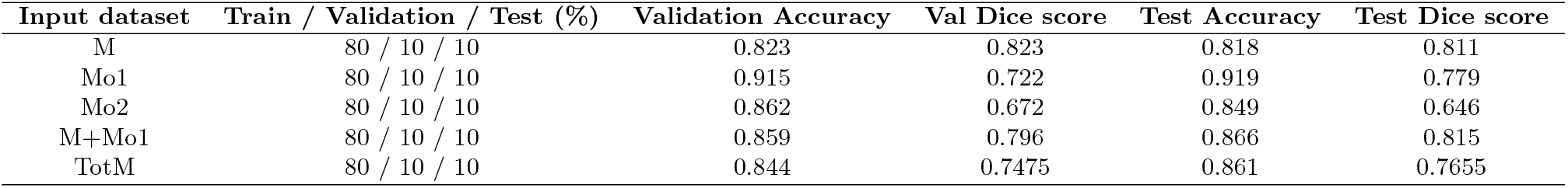
Results on trial with same split train/test but different datasets. The table reports the results obtained for each experiment conducted on the different datasets, including the evaluation metrics computed on both the validation and test sets.

| Input dataset | Train / Validation / Test (%) | Validation Accuracy | Val Dice score | Test Accuracy | Test Dice score |
| --- | --- | --- | --- | --- | --- |
| M | 80 / 10 / 10 | 0.823 | 0.823 | 0.818 | 0.811 |
| Mo1 | 80 / 10 / 10 | 0.915 | 0.722 | 0.919 | 0.779 |
| Mo2 | 80 / 10 / 10 | 0.862 | 0.672 | 0.849 | 0.646 |
| M+Mo1 | 80 / 10 / 10 | 0.859 | 0.796 | 0.866 | 0.815 |
| TotM | 80 / 10 / 10 | 0.844 | 0.7475 | 0.861 | 0.7655 |

**Table 4.** Results on trial with different train/test splits, but same dataset. The table reports the results obtained across different experiments with varying train/test split ratios for the M+Mo1 dataset.

| Input dataset | Train / Validation / Test (%) | Validation Accuracy | Val Dice score | Test Accuracy | Test Dice score |
| --- | --- | --- | --- | --- | --- |
| M+Mo1 | 80 / 10 / 10 | 0.859 | 0.796 | 0.866 | 0.815 |
| M+Mo1 | 60 / 10 / 30 | 0.8585 | 0.815 | 0.874 | 0.819 |
| M+Mo1 | 40 / 10 / 50 | 0.864 | 0.806 | 0.869 | 0.819 |
| M+Mo1 | 20 / 10 / 70 | 0.8715 | 0.816 | 0.857 | 0.7995 |

### 3.3 Model performance using smaller training datasets

This section aims to explore the capability of self-supervised learning approaches to effectively learn from a reduced amount of annotated data compared to conventional machine learning and deep learning techniques. Typically, the training of such models requires thousands of samples; however, in this study, meaningful results were achieved using only a few hundred training instances. Accordingly, the analysis was designed to investigate how far the proportion of training data could be reduced relative to the total available dataset. To this end, a series of tests was conducted on selected datasets and on combinations thereof, focusing on those that had demonstrated the best performance in the previous training phase. The evaluation aimed to determine whether satisfactory results could still be achieved, in terms of both quantitative metrics and qualitative outputs, when the same dataset was partitioned into training, validation, and testing subsets with a fixed proportion of 80/10/10.

The training parameters were kept consistent with those previously identified as optimal, namely a batch size of 32 and a learning rate of 10^*−*4^. The results reported in Tab. 3 include validation metrics used to assess the training phase and test metrics used to evaluate the model performance on unseen data. Overall, all datasets yielded satisfactory results in terms of evaluation metrics, with the exception of Mo2, which exhibited lower performance. This behaviour can be attributed to the increased complexity of the corresponding masks, as discussed in the previous sections, and can be observed in Fig. 6 and in the images provided in the SI.

For datasets obtained by combining multiple sources, particular attention was paid to preserving the relative proportions of each constituent subset across the training, validation, and testing splits, ensuring consistency with the original dataset distribution.

The analysis was therefore extended by evaluating, for the dataset that achieved the best performance during the testing phase, different proportions between the training and test sets. The results reported in Tab. 4 indicate that the model is not significantly affected by a substantial reduction in the size of the training dataset. This highlights the robustness of the adopted Barlow Twins approach, demonstrating its ability to effectively learn and reproduce relevant features even when only a limited amount of annotated data is available. Notably, high values of accuracy and Dice score are maintained during the testing phase even when only 30% of the data are annotated. It remains essential to complement quantitative evaluation with a careful inspection of the output masks. Although the evaluation metrics appear relatively stable across different training set sizes, qualitative analysis provides additional insight into model performance. In particular, the model does not exhibit difficulties in generating accurate wrinkle masks for data derived from M, whereas it shows greater limitations when applied to data from Mo1. This behaviour can be attributed to the low contrast between wrinkles and background in these images, which makes the segmentation task inherently more challenging, as can be observed in the images provided in the SI.

### 3.4 Sholl analysis

To move beyond a purely qualitative description of biofilm morphology, a quantitative approach was required. Therefore, Sholl analysis was performed to characterize biofilm structure through the spatial distribution and organization of wrinkle patterns. It is important to carefully select the datasets on which the Sholl analysis is performed. In this work, the analysis was primarily explored and discussed on the M dataset, due to limitations of the other datasets.

The masks from the M, Mo1, and Mo2 datasets were selected for a preliminary assessment aimed at determining which datasets were most suitable for Sholl analysis. Representative masks are shown in Fig. 10. As can be observed, the Mo1 dataset contains wrinkles with a predominantly linear and radial morphology, characteristic of *E. coli* AR3110, which limits the amount of structural complexity that can be captured through this type of analysis. In addition, the wrinkles are less clearly distinguishable from the surrounding biofilm background, introducing greater uncertainty during the skeleton extraction step required for Sholl analysis. This issue is particularly evident in the bottom image of the second column. The C1 dataset originates from the same group of *E. coli* biofilms and therefore presents similar limitations to Mo1. By contrast, the masks from the Mo2 dataset, and consequently those from C2, as both datasets originate from the same experiment on *B. subtilis*, exhibit wrinkle patterns characterized by a less circular geometry. Furthermore, the wrinkles are predominantly concentrated in the peripheral region of the colony, whereas the central area remains largely empty and provides little structural information, as shown in the third column of Fig. 10, and characteristic of *B. subtilis* biofilms grown in non-ideal concentrations of glutamate and glycerol. These morphological characteristics make the M dataset the most suitable for Sholl analysis. Its wrinkles are clearly distinguishable from the remainder of the biofilm and form an approximately circular architecture that is well suited to an analysis based on concentric circles. The masks generated by the algorithm during the testing phase were also considered too preliminary for this purpose, as they did not provide sufficient accuracy and remained affected by noise. Reliable Sholl analysis capable of yielding meaningful quantitative information on wrinkle organization requires masks that are both clean and accurately representative of the underlying biofilm structure.

**Fig. 10.**
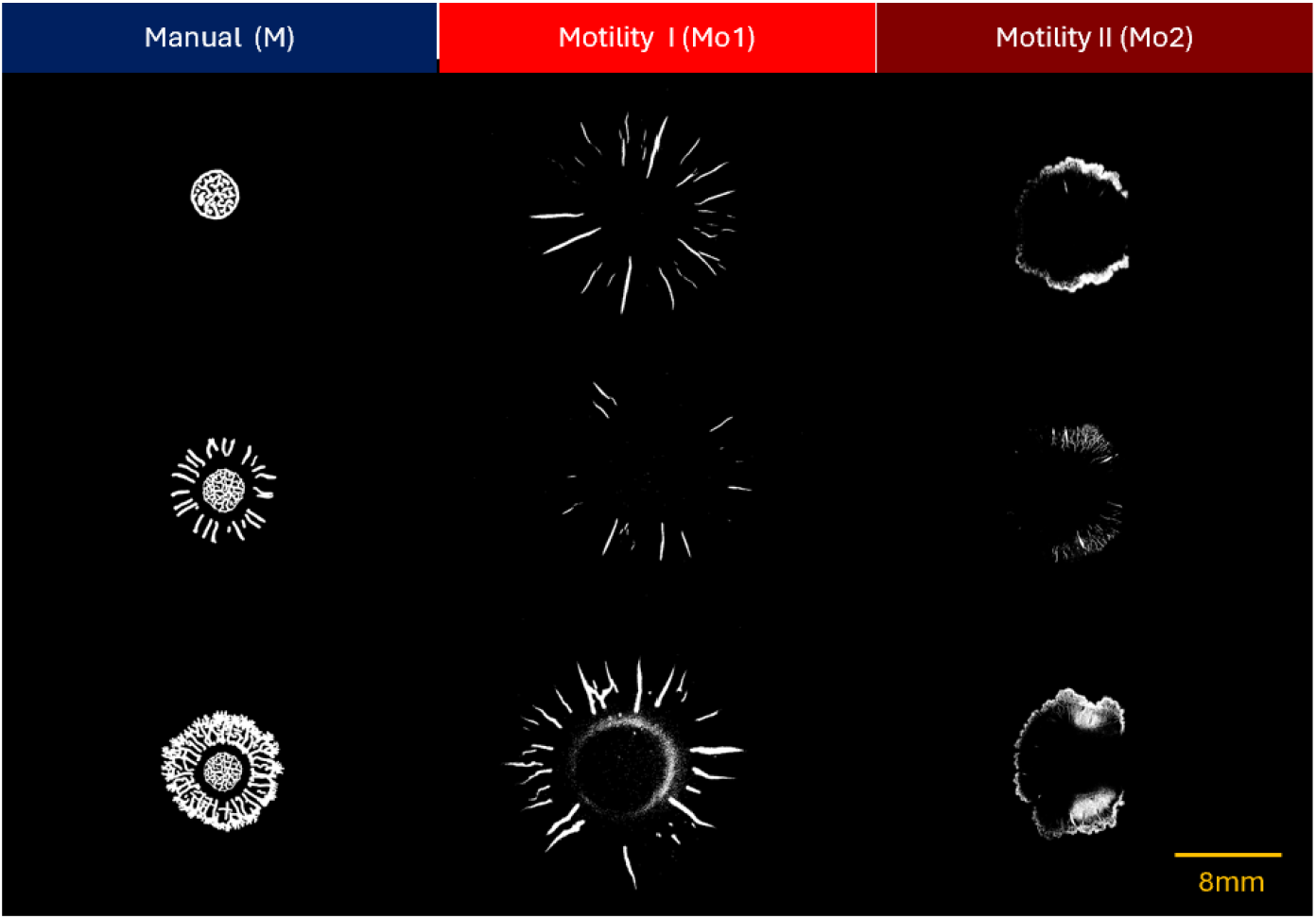
Representative masks from the M, Mo1, and Mo2 datasets. The first column shows several examples of masks from the M dataset. The wrinkles are clearly distinguishable from the remainder of the biofilm and exhibit an approximately circular and highly branched architecture, making them well suited for Sholl analysis. The second column shows masks from the Mo1 dataset, in which the wrinkles display a predominantly linear morphology. In the bottom image, noise unrelated to wrinkle structures is also visible in the central region of the biofilm. The third column presents masks from the Mo2 dataset, characterized by a noncircular colony morphology and by wrinkles mainly concentrated in the peripheral region of the biofilm. In addition, the boundaries and shapes of the wrinkles are less clearly defined.

The dataset M was constructed from 12 time-lapse sequences representing the evolution of *B. subtilis* biofilms of strain NCIB 3610 and comprises a total of 58 images, corresponding to 58 annotated masks. The 12 time-lapse sequences, spanning from frame 1 to 175, the time interval between the acquisition of two consecutive frames was 40 minutes, corresponding to a total duration of nearly 117 hours, or approximately 5 days. It is of particular interest to investigate the Sholl analysis across different stages of biofilm development in order to highlight structural differences through parameters that go beyond simple size measurements. The initial implementation of the code was adapted from a publicly available repository on GitHub [18].

The Sholl analysis reveals that the evolution of this set of biofilm, *B. subtilis* of strain NCIB 3610 grown on MSgg media, can be clearly divided into three distinct phases. Through Sholl analysis, these three phases can be quantitatively characterized in a more rigorous manner, first within a single temporal sequence and subsequently across all available datasets. Accordingly, the analysis of one representative biofilm is presented across 8 images describing its temporal evolution in Fig. 11, and is used to define the parameters later applied to the complete dataset.

**Fig. 11.**
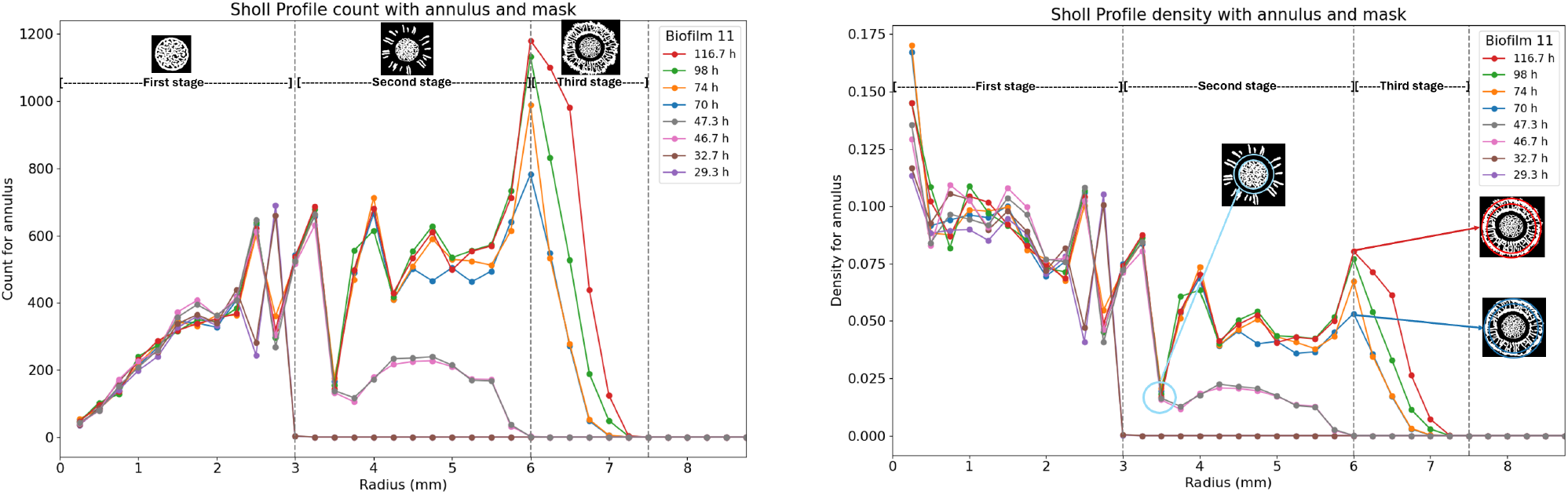
Tracking biofilm evolution for one biofilm using Sholl analysis. a) the number of skeleton pixels counted within each annulus is shown, alongside b) the corresponding density per annulus. The analysis is performed across the 8 available images belonging to the same time lapse sequence. The labels reported in the legend correspond to the frame number of each image within the time lapse. This analysis produces a quantitative description of the structural evolution of the biofilm over time and helps explain the bimodal distribution of biofilm radii observed in the M dataset. In particular, the Sholl profiles reveal that the evolution of the biofilm is not characterized by two stages, but rather by three distinct phases. The analysis of this single biofilm was therefore used as a reference to extend the study to the entire M dataset, which was subsequently divided into three subsets corresponding to the three evolutionary stages highlighted here. The first phase, in which the biofilm diameter does not exceed approximately 3 mm, extends up to frame 50 of the time lapse sequence. The second phase is characterized by continued radial growth together with the progressive formation of wrinkles, reaching approximately 6 mm, and extends up to frame 100. The third and final phase exhibits a high wrinkle density, with biofilm dimensions reaching approximately 7.25 mm and a more complex structural organization. An additional observation can be drawn from the density plot. In the final stages of evolution, the wrinkle density in the outer region of the biofilm reaches values comparable to those observed in the more complex inner region. This occurs despite the presence of a pronounced gap around 3.5 mm, where wrinkles are largely absent for all biofilms that extend beyond this radius. This behaviour suggests that the biofilm does not develop wrinkles in this intermediate region, while still continuing to expand and form new wrinkles in the outermost area.

Based on this analysis, three distinct developmental phases can be identified, which are clearly distinguishable in the graph showing the wrinkle counts for each annulus. The first phase extends up to frame 50 of the time lapse sequence. During this stage, the biofilm does not exceed approximately 3 mm in radial extent, while exhibiting a high wrinkle density. The second phase extends to frame 100 and is characterized by continued biofilm growth and the progressive formation of wrinkles up to a radial distance of approximately 5 mm. In addition, a region around 3.5 mm displays a particularly low wrinkle density per annulus, corresponding to an internal area of the biofilm in which wrinkles are largely absent. During the third and final phase, the wrinkles continue to develop and extend to approximately 7.25 mm. At this stage, the wrinkle density per annulus increases again, reaching values comparable to those observed in the innermost region of the biofilm. Since the smallest biofilm has a diameter of approximately 6 mm, corresponding to a radius of approximately 3 mm or 120 pixels, an estimated spatial conversion of approximately 0.025 mm per pixel can be obtained.

The analysis of branching points and endpoints, shown in Fig. 12, indicates that in the final images of the third phase the biofilm undergoes further structural development while maintaining approximately the same overall diameter. Notably, the increase in both branching points and endpoints suggests that, rather than expanding outward, the wrinkle network within the biofilm becomes increasingly fragmented and spreads throughout regions that are already occupied by wrinkles.

**Fig. 12.**
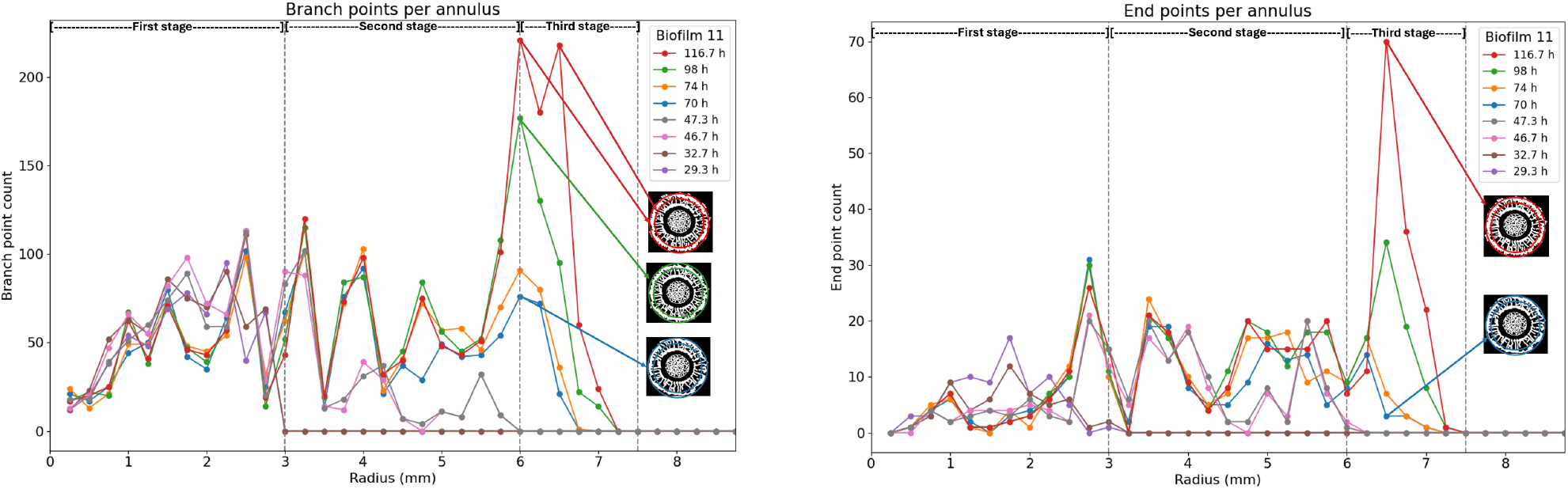
Tracking biofilm complexity for one biofilm using Sholl analysis. The branching points analysis is displayed on the left and endpoints in right panel. This more detailed analysis reinforces the observations discussed in the previous section. In particular, the branching point profile exhibits strong variability in the region between approximately 5.5 and 6.75 mm. This behaviour suggests that, rather than extending further in radius, the biofilm continues to increase its structural complexity by generating additional wrinkles within an already explored region. The endpoint analysis shows a relatively regular trend across most stages, with the exception of the curve corresponding to the final stage of biofilm evolution. In this case, a pronounced increase in endpoints is observed in the outermost region, indicating the formation of newly developing wrinkle tips at the biofilm periphery.

The analysis was conducted separately for each stage, and in Fig. 13 each plot shows the mean values, along with the corresponding standard deviation, for the three evolutionary stages. This study confirms that the division of biofilm evolution into three phases is well supported. In the region between 5 and 6.25 mm, corresponding to the third phase, a high standard deviation is observed compared to the rest of the graph, indicating that while different biofilms exhibit similar behaviour during the initial stages, their trajectories diverge in the later stages of development.

**Fig. 13.**
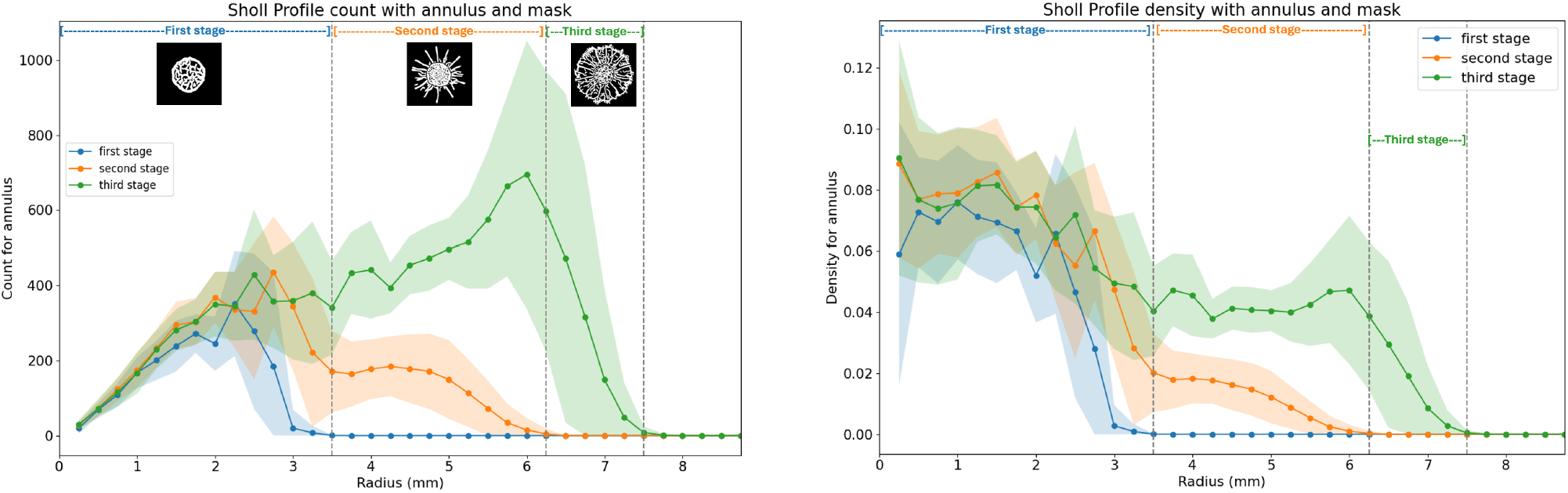
Tracking biofilm evolution for all biofilms in M dataset using Sholl analysis. a) wrinkle counts and b) wrinkle density. a) the number of counts per annulus is reported, alongside b) the density per annulus, both computed for the three phases of biofilm evolution. The solid line represents the mean, whereas the shaded area indicates the standard deviation. This analysis builds upon the observations derived from the temporal evolution of a single biofilm. The obtained values are consistent with those observed in the single biofilm case, supporting the validity of the three phase division for this dataset and highlighting the structural complexity and variability of the central regions characteristic of this biofilm morphology. In the count plot, it is particularly interesting to observe the large standard deviation in the outer region of the third evolutionary stage. This behaviour suggests that, rather than continuing to expand radially, the biofilm tends to increase structural complexity within regions that have already been explored. The relatively low standard deviation in the central region suggests that this area is fairly homogeneous across all biofilms in which it is present. In contrast, the density plot shows a higher standard deviation during the early stages of evolution, indicating greater variability in wrinkle formation when the biofilm structure is still developing.

A similar trend is observed in the analysis of branching points, endpoints, and internal points Fig. 14, where the overall behaviour remains consistent but a greater variability emerges in the final values of the third phase.

**Fig. 14.**
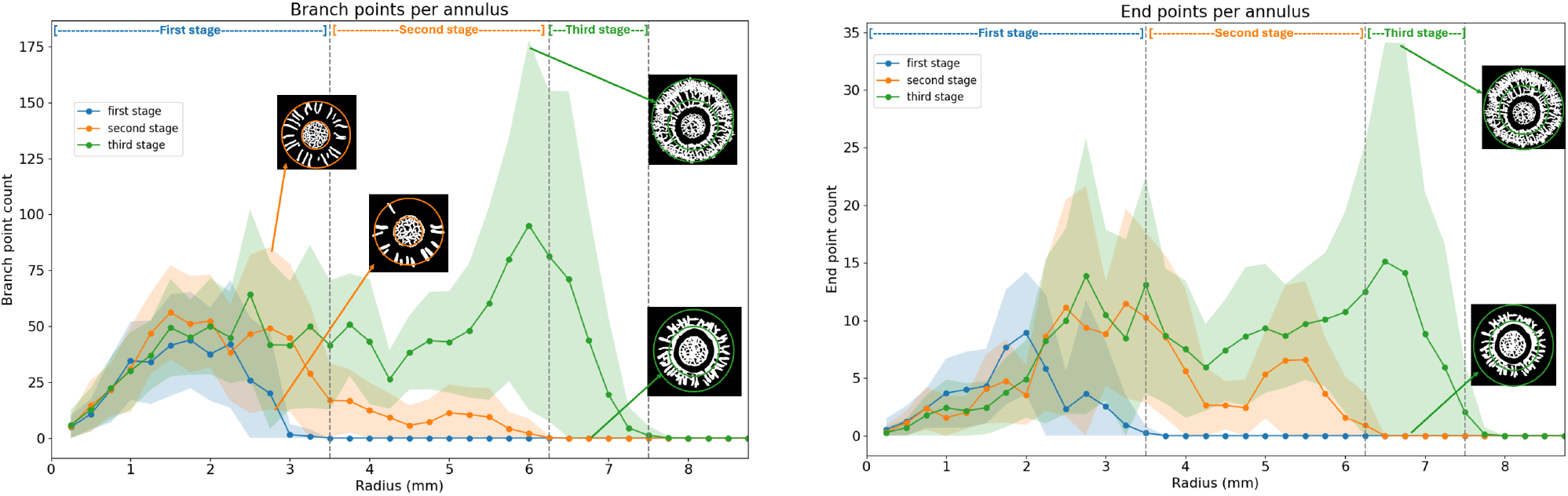
Tracking biofilm complexity for all biofilms in M dataset using Sholl analysis. The branching points analysis is displayed on the left and endpoints in right panel. This analysis once again confirms the validity of the three phase division for this dataset. The overall trends are consistent with those observed for the single biofilm, while also highlighting the variability across different samples. The greatest variability is observed in the region between approximately 5 and 6.25 mm. Although the overall trends remain consistent, the large standard deviation indicates that, across different biofilms or across different stages of the same biofilm, the final phase is considerably more heterogeneous. This behaviour is evident in all three plots, and particularly in the branching point profile, where the variability is especially pronounced. This suggests that in this outer region the biofilm no longer primarily expands radially, but instead increases structural complexity by branching within an already explored area. The endpoint analysis shows high variability also in the central region, particularly between 2.5 and 3.75 mm. This indicates that some biofilms exhibit a large number of wrinkle tips even in the central area, making this region highly heterogeneous for both the second and third evolutionary stages.

## 4 Conclusion

In conclusion, this study demonstrates that the proposed pipeline, AutoWrinkleID, is capable of extracting masks containing only the wrinkle structures of biofilms. This approach offers a scalable and automated alternative to manual annotation, as well as to fluorescence-based imaging methods, which require genetic modification of the bacterial strains. It enables the analysis of biofilm morphology across different experimental conditions. The results obtained on the validation sets of several datasets achieved accuracy and Dice score values around or above 80% (Tab. 2),indicating a strong correspondence between the predicted masks and the ground truth. It is important to note that during the training phase the models were able to reconstruct masks that were more accurate than the manual annotations. This behaviour arises because the algorithm learns the structural features directly from the raw biofilm images, while the manual annotations are incorporated only at a later stage as supervisory guidance rather than as the primary source of information. The performance of models trained on masks derived from motility and constitutive fluorescence signals was comparable during the training phase in terms of validation metrics. However, models trained on motility-based masks exhibited improved accuracy and greater flexibility during the testing phase, highlighting their superior generalization capability across the different test datasets. It was further observed that, even with only 30% of annotated data (20% used for training and 10% for validation), the algorithm remains capable of correctly identifying biofilm wrinkles. The application of Sholl analysis allowed for a detailed quantitative characterisation of the M dataset, demonstrating the effectiveness of this approach for the study and analysis of biofilm wrinkles, as it is capable of capturing their structural complexity. However, this method proved to be less informative for datasets in which the wrinkles exhibited less branched morphologies and for images affected by substantial noise, conditions that limit the ability of the analysis to accurately quantify wrinkle. The main limitations of this work include the relatively limited number of biofilms analysed and the quality of the masks used during training, which directly influence the learning process of the algorithm. Future developments will focus on the the inclusion of additional biofilm types in the training data to further improve the flexibility and generalization capability of the algorithm, and the development of advanced analysis methods based on Sholl analysis to link extracted features to specific biofilm types.

## 5 Acknowledgments

The authors would like to thank Dr. Ashraf Zarkan and Kieran Abbott from the Department of Genetics, University of Cambridge, for providing the strains for this study.

## 6 Data Availability

The data that support the findings of this study are available upon request.

## 7 Supporting Information

Supporting Information is available for this study. We present additional evaluation metrics, as well as further model outputs from models trained on Mo1, Mo2, C1 and C2 datasets.

