## Supplementary information file for "AutoWrinkleID: a machine learning pipeline for biofilm wrinkle identification and quantitative analysis"

### Table of Contents

**Supplementary Fig. S1: Evaluation curves for the validation subset during training.** **2**

**Supplementary Tab. S1: Evaluation metrics overview for the validation subsets.** **3**

**Supplementary Fig. S2: Outputs from the model trained on the Mo1 and Mo2 datasets.** **3**

**Supplementary Fig. S3: Outputs from the model trained on the C1 and C2 datasets.** **4**

**Supplementary Fig. S4: Results on Test 1.** **4**

**Supplementary Fig. S5: Results on Test 2.** **5**

**Supplementary Fig. S5: Results on Test 3.** **5**

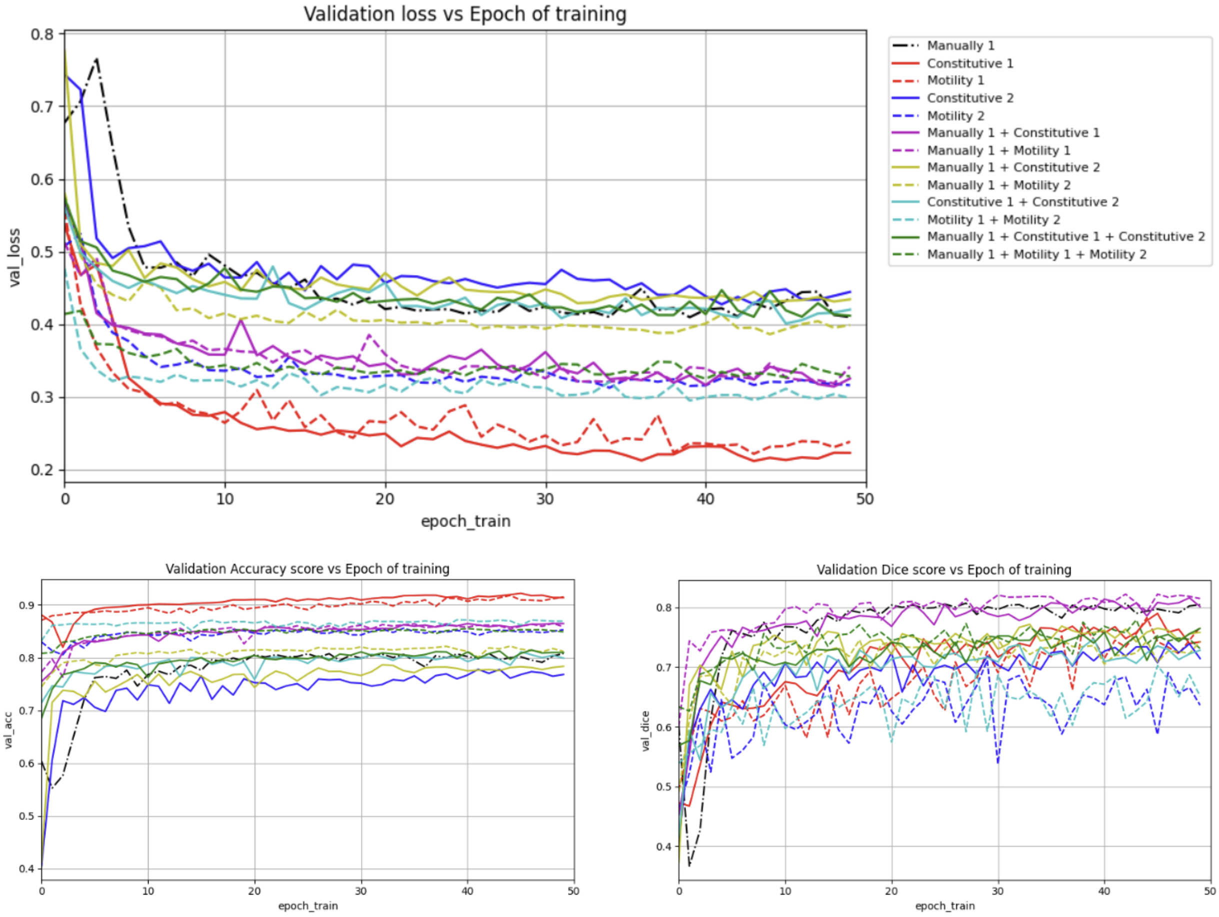


**Fig. 1 Evaluation curves for the validation subset during training.** The top panel shows the loss curves for all training datasets and their combinations. The bottom left panel reports the accuracy, while the bottom right panel shows the Dice score. Results are shown for the validation subsets of each dataset and their combinations. The evolution of these metrics across epochs is as important as the final values reported in Tab. 1, as it demonstrates that the different training configurations do not exhibit overfitting and that convergence is reached by the end of the training phase, which was performed for 50 epochs for all models, they have also the same learning rate of $10^{-4}$. These plots are also essential for selecting which model weights should be retained for the testing phase. Most curves obtained from constitutive fluorescence datasets show trends consistent with their corresponding motility fluorescence counterparts. The training metrics must then be interpreted together with the qualitative outputs of the algorithm presented in the following section.

| Dataset ID | Dataset | Images | Accuracy | Dice score |
|---|---|---|---|---|
| M | Manual | 180 | 0.81 | **0.80** |
| C1 | Constitutive I | 150 | 0.91 | 0.74 |
| Mo1 | Motility I | 260 | **0.915** | 0.77 |
| C2 | Constitutive II | 340 | 0.77 | 0.71 |
| Mo2 | Motility II | 340 | 0.85 | 0.64 |
| M+C1 | Manual + Constitutive I | 330 | 0.865 | 0.80 |
| M+Mo1 | Manual + Motility I | 440 | 0.86 | **0.815** |
| M+C2 | Manual + Constitutive II | 520 | 0.78 | 0.76 |
| M+Mo2 | Manual + Motility II | 520 | 0.81 | 0.74 |
| C1+C2 | Constitutive I + Constitutive II | 490 | 0.80 | 0.73 |
| Mo1+Mo2 | Motility I + Motility II | 600 | **0.87** | 0.65 |
| M+C1+C2 | Manual + Constitutive I + Constitutive II (Total Constitutive) | 670 | 0.81 | **0.76** |
| M+Mo1+Mo2 | Manual + Motility I + Motility II (Total Motility) | 780 | **0.85** | 0.73 |

**Table 1 Evaluation metrics overview for the validation subsets.** Overview of the datasets used for training, including the number of image patches and the evaluation metrics obtained for the corresponding validation subsets. All models were trained for 50 epochs using a learning rate of $10^{-4}$ and a batch size of 32. Results are reported for the validation subsets of the individual datasets and their combinations.

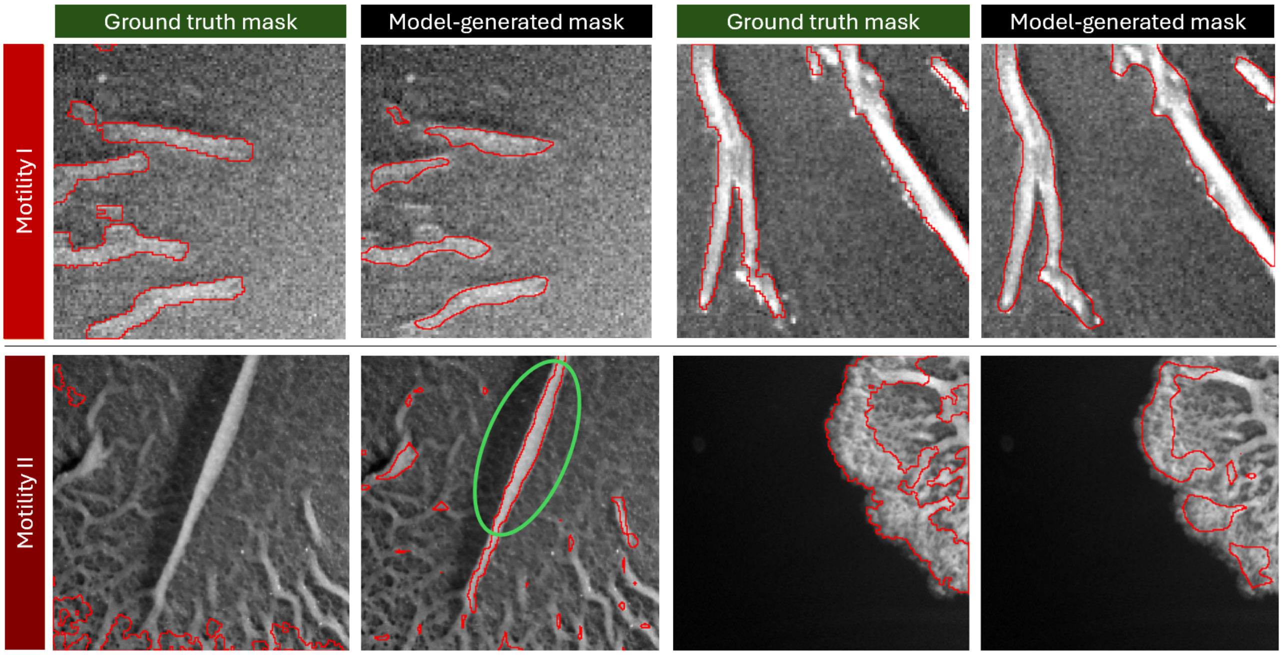


**Fig. 2 Outputs from the model trained on the Mo1 and Mo2 datasets.** These examples illustrate the performance of the model when trained on these datasets. The green circle in the second panel of the first row highlights an instance in which the model identifies the complexity of the wrinkle pattern more accurately than the corresponding motility fluorescence annotation. For the Mo1 dataset, the limited contrast between the wrinkles and the surrounding biofilm affects the quality of both the masks generated from motility related fluorescence and those predicted by the model trained on these data. In the Mo2 dataset, by contrast, the high morphological complexity of the wrinkle patterns makes it difficult to generate masks that accurately delineate the wrinkles.

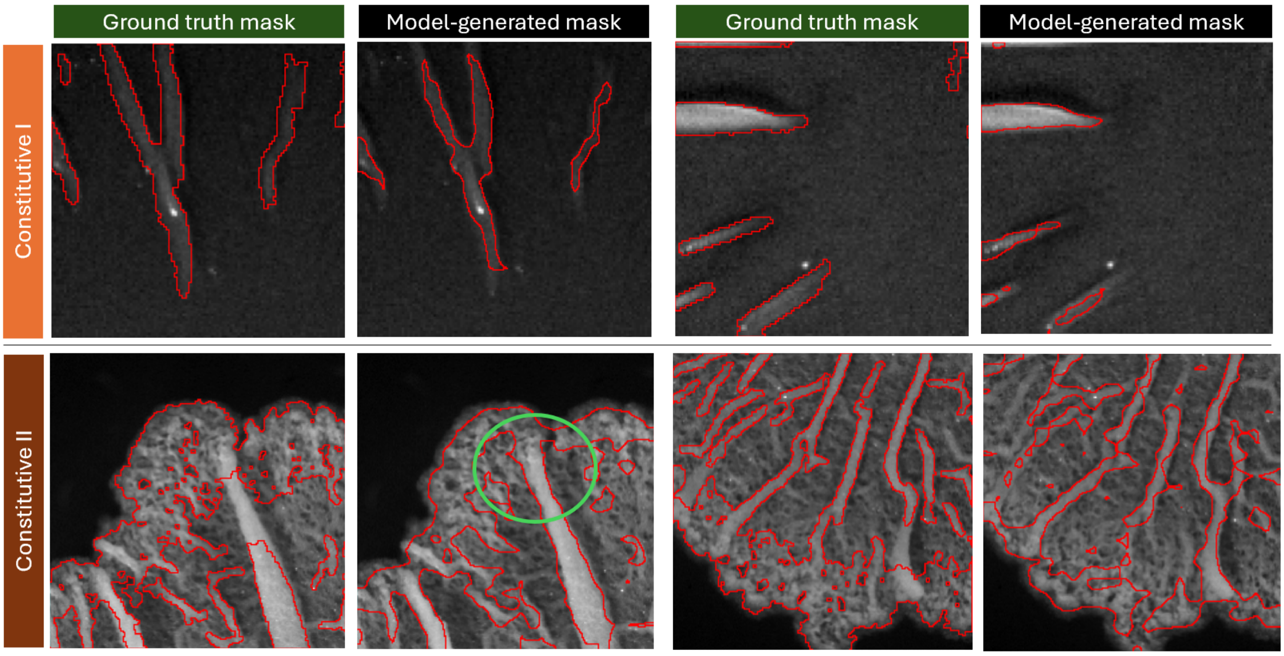


**Fig. 3 Outputs from the model trained on the C1 and C2 datasets.** These examples illustrate the performance of the model when trained on these datasets. The green circle in the second panel of the first row highlights an instance in which the model identifies the complexity of the wrinkle pattern more accurately than the corresponding constitutive fluorescence annotation. For this training dataset, the model identifies the wrinkles more accurately than the model trained on the combined Mo1 and Mo2 datasets. Although the Mo1+Mo2 model achieves a higher accuracy, the model trained on this dataset obtains a higher Dice score, indicating a better overlap between the predicted masks and the ground truth annotations. Constitutive fluorescence provides a clearer representation of the wrinkles in the C2 subset, thereby enabling the model to identify and delineate the wrinkle structures more accurately.

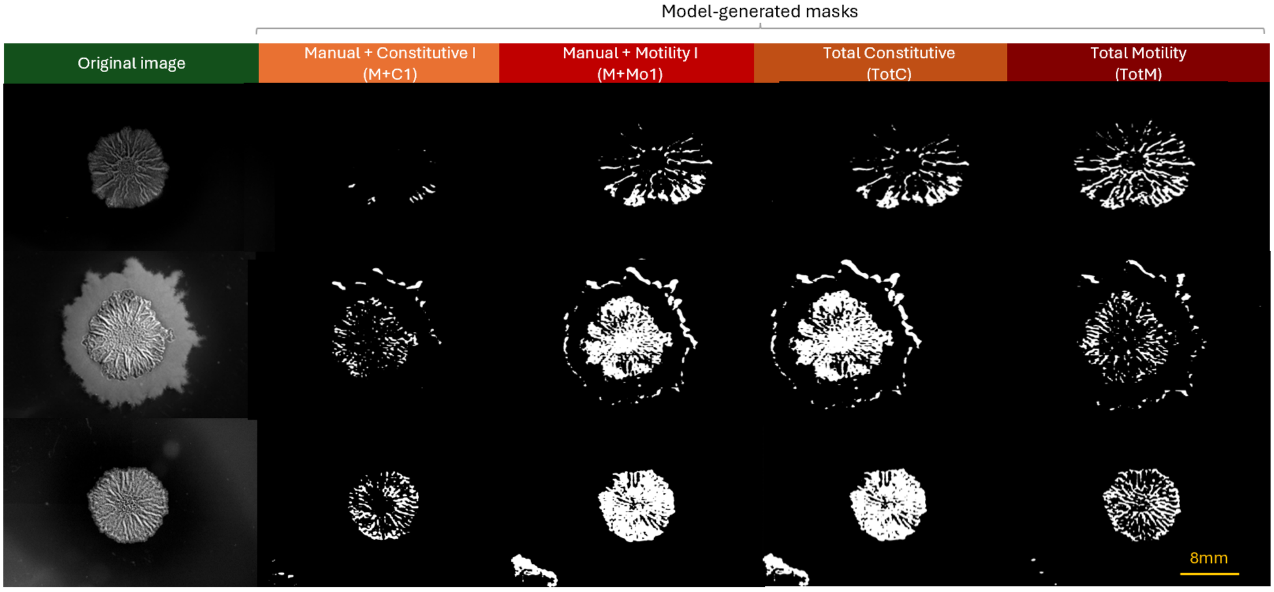


**Fig. 4 Results on Test 1.** The data in this dataset, referred to as Test 1, are similar to those in the M dataset. Examples are presented in which the algorithm fails to identify the wrinkles accurately.

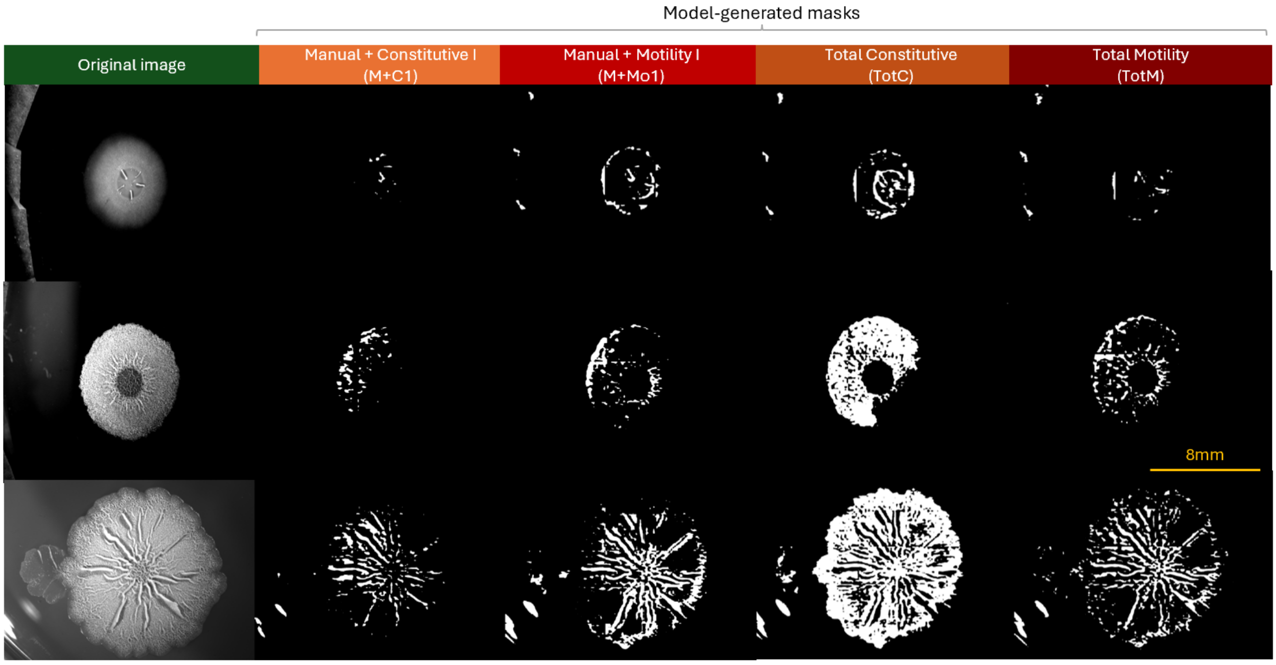


**Fig. 5** **Results on Test 2.** The data in this dataset , referred to as Test 2, are similar to those in the C2 and Mo2 datasets. In this section, examples are presented in which the algorithm was unable to identify the wrinkles accurately. The results also show that the algorithm is sensitive to artefacts present in the images.

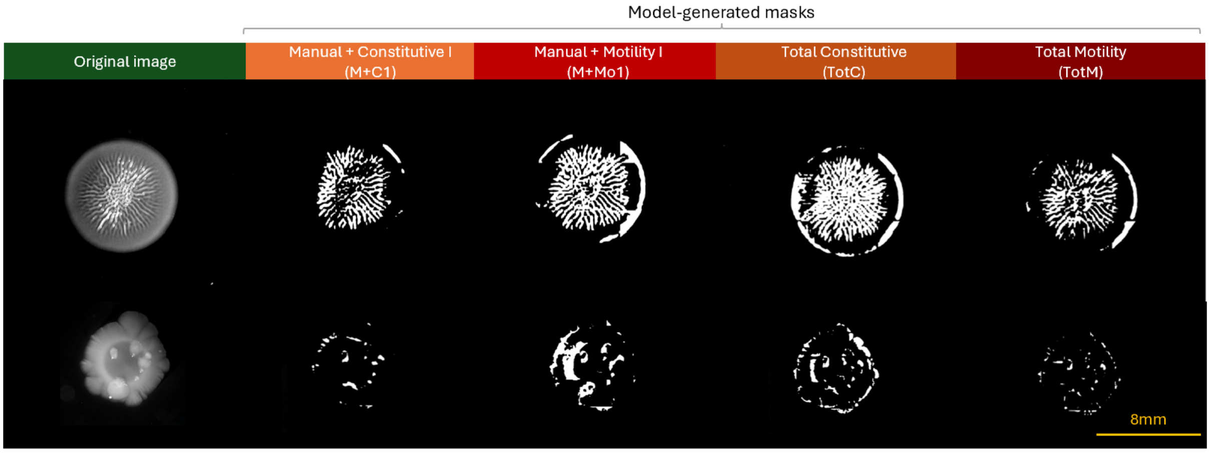


**Fig. 6** **Results on Test 3.** This test dataset, referred to as Test 3, represents a challenging scenario for the model, as the data differ significantly from those used during training. In this case, examples were selected in which the algorithm encountered difficulties in correctly identifying the wrinkles.
